# A CRISPR interference platform for selective downregulation of gene expression in *Borrelia burgdorferi*

**DOI:** 10.1101/2020.10.12.335471

**Authors:** Constantin N. Takacs, Molly Scott, Yunjie Chang, Zachary A. Kloos, Irnov Irnov, Patricia A. Rosa, Jun Liu, Christine Jacobs-Wagner

**Affiliations:** Department of Biology and ChEM-H Institute, Stanford University, Stanford, CA, 94305, USA; Microbial Sciences Institute, Yale West Campus, West Haven, CT, 06516, USA; Department of Molecular, Cellular, and Developmental Biology, Yale University, New Haven, CT, 06511, USA; Howard Hughes Medical Institute, Stanford University, CA, 94305, USA; Department of Microbial Pathogenesis, Yale University School of Medicine, New Haven, CT, 06511, USA; Microbiology Program, Yale University, New Haven, CT, 06511, USA; Laboratory of Bacteriology, Rocky Mountain Laboratories, Division of Intramural Research, National Institute of Allergy and Infectious Diseases, National Institutes of Health, Hamilton, MT, 59840, USA

**Keywords:** CRISPR, *Borrelia*, Lyme disease, spirochete, dCas9, cell morphogenesis, bacteria, MreB, RodA, FtsI

## Abstract

The spirochete *Borrelia burgdorferi* causes Lyme disease, an increasingly prevalent infection. While previous studies have provided important insight into *B. burgdorferi* biology, many aspects, including basic cellular processes, remain underexplored. To help speed up the discovery process, we adapted a CRISPR interference (CRISPRi) platform for use in *B. burgdorferi*. For efficiency and flexibility of use, we generated various CRISPRi template constructs that produce different basal and induced levels of *dcas9* and carry different antibiotic resistance markers. We characterized the effectiveness of our CRISPRi platform by targeting the motility and cell morphogenesis genes *flaB, mreB, rodA,* and *ftsI,* whose native expression levels span two orders of magnitude. For all four genes, we obtained gene repression efficiencies of at least 95%. We showed by darkfield microscopy and cryo-electron tomography that flagellin (FlaB) depletion reduced the length and number of periplasmic flagella, which impaired cellular motility and resulted in cell straightening. Depletion of FtsI caused cell filamentation, implicating this protein in cell division in *B. burgdorferi*. Finally, localized cell bulging in MreB- and RodA-depleted cells matched the locations of new peptidoglycan insertion specific to spirochetes of the *Borrelia* genus. These results therefore implicate MreB and RodA in the particular mode of cell wall elongation of these bacteria. Collectively, our results demonstrate the efficiency and ease of use of our *B. burgdorferi* CRISPRi platform, which should facilitate future genetic studies of this important pathogen.

**IMPORTANCE:** Gene function studies are facilitated by the availability of rapid and easy-to-use genetic tools. Homologous recombination-based methods traditionally used to genetically investigate gene function remain cumbersome to perform in *B. burgdorferi*, as they often are relatively inefficient. In comparison, our CRISPRi platform offers an easy and fast method to implement as it only requires a single plasmid transformation step and IPTG addition to obtain potent (>95%) downregulation of gene expression. To facilitate studies of various genes in wild-type and genetically modified strains, we provide over 30 CRISPRi plasmids that produce distinct levels of *dcas9* expression and carry different antibiotic resistance markers. Our CRISPRi platform represents a useful and efficient complement to traditional genetic and chemical methods to study gene function in *B. burgdorferi*.

## INTRODUCTION

*Borrelia burgdorferi*, a spirochetal bacterium, is maintained in nature via a transmission cycle between a tick vector and a warm-blooded host, such as a wild mouse (1). A bite by a *B. burgdorferi*-colonized tick can lead to transmission of the bacterium to humans. In the absence of timely antibiotic treatment, infection by *B. burgdorferi* causes Lyme disease, a prevalent vector-borne disease in temperate regions of the Northern hemisphere (2). The infection can result in a wide range of symptoms, from malaise, fever, and a characteristic skin rash during early stages of the disease, to cardiac, neurologic, or articular pathologies in later stages (3). The rapid rise in Lyme disease incidence in recent years (2) underscores the need for a detailed understanding of not only the disease, but also the pathogen itself.

Studying the biology of a pathogen is greatly facilitated by the genetic tractability of the organism. For this reason, a diverse set of genetic reagents and techniques have been developed and validated for use in *B. burgdorferi* (4–6). However, creating gene deletion mutants in *B. burgdorferi* remains tedious and time-consuming. Homologous recombination of suicide vectors is needed to create gene deletion mutants. In *B. burgdorferi,* this process occurs at frequencies of ∼10^−7^ (7–9), which are close to the 10^−9^ to 10^−7^ range of frequencies at which spontaneous mutants conferring resistance to common selection antibiotics arise (10). This inefficient process is compounded by *B. burgdorferi*’s slow in vitro growth rate, with typical doubling times in the range of 5 to 18 hours (11–16). Genetic modification is particularly challenging for genes essential for viability; such genes cannot be deleted, and generation of conditional mutants (17) often requires multiple transformation steps and is therefore even slower and less efficient.

The development of CRISPR (clustered regularly interspaced palindromic repeats) interference (18, 19), or CRISPRi, provides an attractive complement to traditional genetic manipulation protocols based on homologous recombination. CRISPRi is a gene product depletion method that requires two components. One of them is dCas9, a catalytically inactive version of Cas9, which is the nuclease component of a bacterial adaptive immunity system against invading DNA molecules (18, 20, 21). The other component is a short guide RNA molecule, or sgRNA. The sgRNA (Fig. 1A) contains a base-pairing region and a dCas9 recognition loop, which is called a dCas9 handle (18, 20). The base-paring region is usually a 20-nucleotide stretch complementary to the target DNA strand (18, 19). Proper targeting also requires a protospacer adjacent motif, or PAM, located in the DNA next to the base-pairing region. When co-expressed, dCas9 and the sgRNA form a complex that scans the DNA until it finds a PAM-proximal sequence complementary to the sgRNA’s base-pairing region (22). There, the dCas9-sgRNA complex stably binds to the DNA. When dCas9 is targeted to a promoter or operator, it blocks transcription initiation. When dCas9 is targeted to the coding sequence or the 5′ untranslated region (5′ UTR) of a gene, it blocks transcription elongation (18). Together, the required sgRNA-DNA complementary base pairing and presence of an adjacent PAM site render CRISPRi highly specific (18). Since its development, the CRISPRi method has been adapted for the study of a wide variety of organisms and cell types, from mammalian cells and yeast (23) to various bacterial species, including *E. coli* (18), *Bacillus subtilis* (24, 25), *Caulobacter crescentus* (26, 27), *Mycobacterium tuberculosis* (28–30), and *Leptospira interrogans* (31) to name just a few.

**Figure 1.**
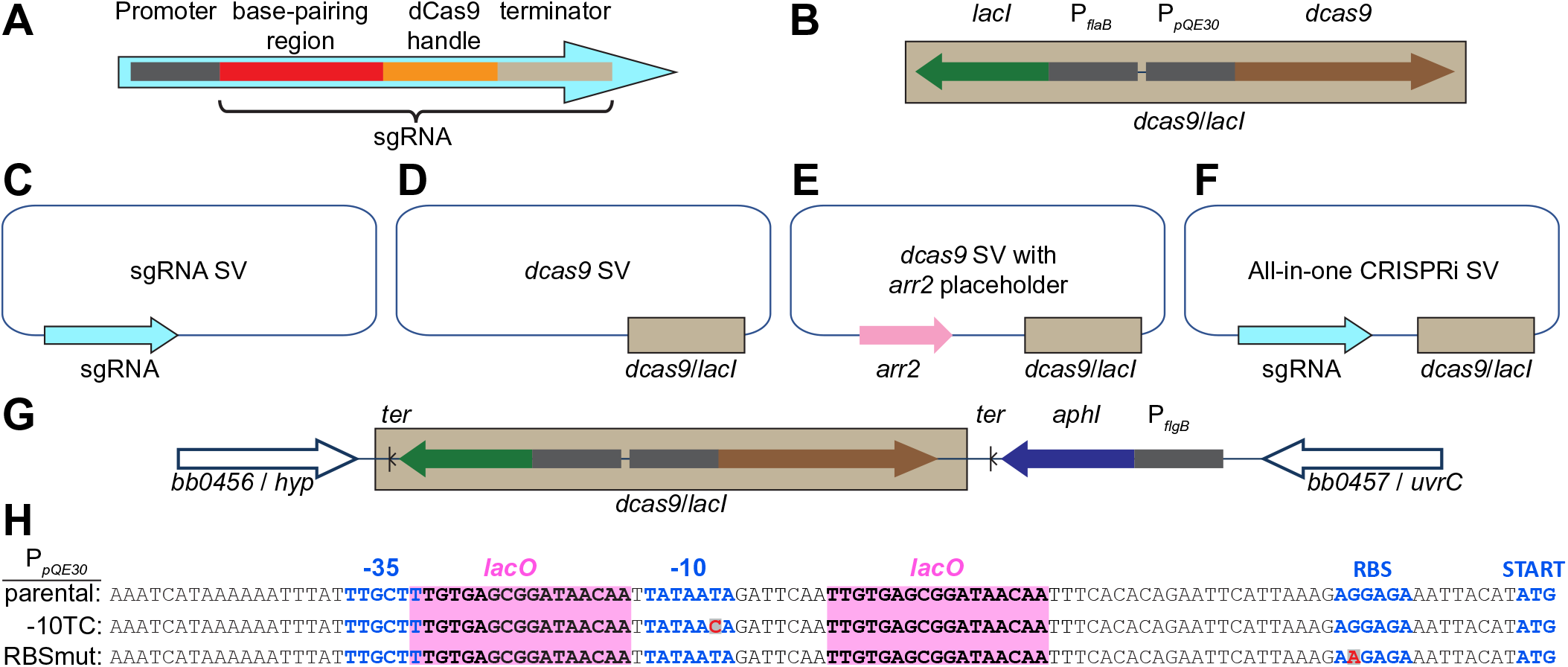
Summary of relevant genetic constructs. **A.** Schematic of a mature sgRNA cassette, including its promoter and relevant functional parts. **B.** Schematic of the inducible *dcas9/lacI* cassette. **C-F.** Schematic of shuttle vectors (SV) generated and used in this study. The *B. burgdorferi* and *E. coli* origins of replication and the antibiotic resistance markers are not shown. *arr2*, rifampin resistance cassette. **G.** Schematic of the chromosomally encoded *dcas9* locus of strain CJW_Bb362. The *dcas9/lacI* cassette and a kanamycin resistance cassette were inserted in the intergenic region between genes *bb0456* and *bb0457*. *ter*, transcriptional terminator; *aphI*, kanamycin resistance gene. **H.** Mutations introduced into the P*_pQE30_* sequence that successfully decreased basal expression of *dcas9*. *lacO*, LacI binding sites; RBS, ribosomal binding site. Mutated residues are shown in red.

Here, we have adapted the CRISPRi system for use in *B. burgdorferi*. We present multiple versions of the platform that are characterized by efficient gene product downregulation, ease of use, and relatively fast clone generation. We have evaluated the functionality of these versions by targeting genes that have broadly varied native expression levels and are involved in *B. burgdorferi* motility, cell shape determination, and cell division.

## RESULTS

### Constructs and strains for sgRNA and *dcas9* expression in *B. burgdorferi*

To adapt the CRISPRi system for use in *B. burgdorferi*, we assembled expression cassettes for sgRNAs and *dcas9* (Fig. 1A-B). The mature sgRNA cassettes that we designed contain the sgRNA sequence fused to the transcriptional start site of a constitutive promoter (Fig. 1A). To generate mature sgRNA cassettes, we first assembled “template” cassettes, which contain the promoter driving sgRNA expression, a ∼500 base pair DNA filler sequence derived from a firefly luciferase gene (32) and the sgRNA’s dCas9 handle (Fig. S1A). We then released the DNA filler sequence by digestion with SapI (or its isoschiomers BspQI or LguRI) and ligated in its place a pair of annealed primers (Fig. S1A-B). We designed these primers such that they encode the sgRNA’s base-pairing region (see Materials and Methods). For sgRNA expression, we primarily employed a synthetic promoter, J23119, hereafter referred to as P*_syn_*, which was previously used to drive sgRNA expression in *E. coli* (18, 19). Using a transcriptional fusion to mCherry, we showed that P_syn_ is active in *B. burgdorferi* (Fig. S1C-D). To drive sgRNA expression, we also tested multiple *B. burgdorferi* promoters (Table S1) whose strengths we previously characterized (Fig. S1D) (6). The nature of the promoter driving sgRNA expression, however, did not appear to affect the functionality of the CRISPRi platform (see below). We generated both the template and the mature sgRNA cassettes in *B. burgdorferi/E. coli* shuttle vectors and refer to them as sgRNA shuttle vectors (Fig. 1C and S1A and Table 1).

**Table 1.**
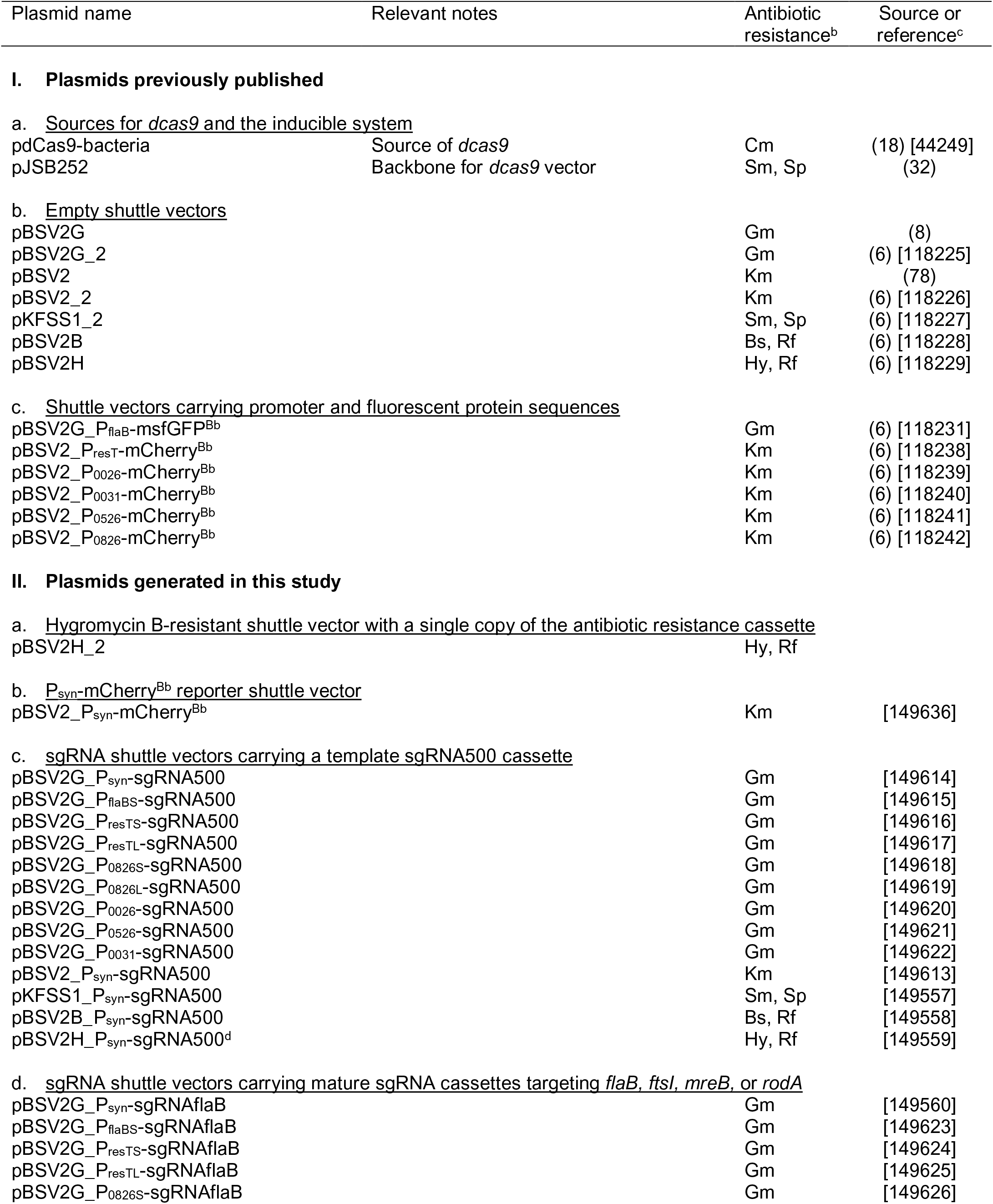
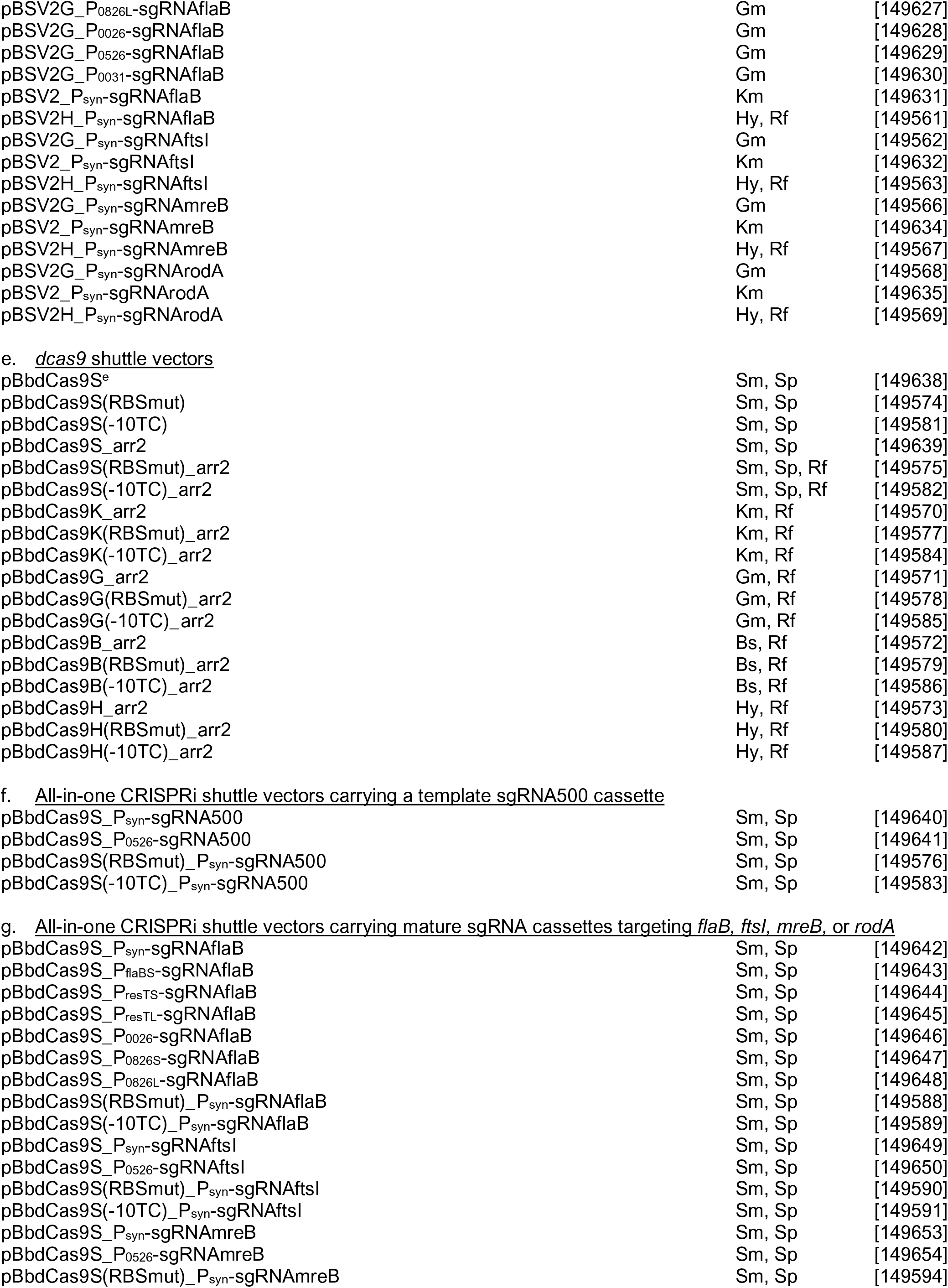
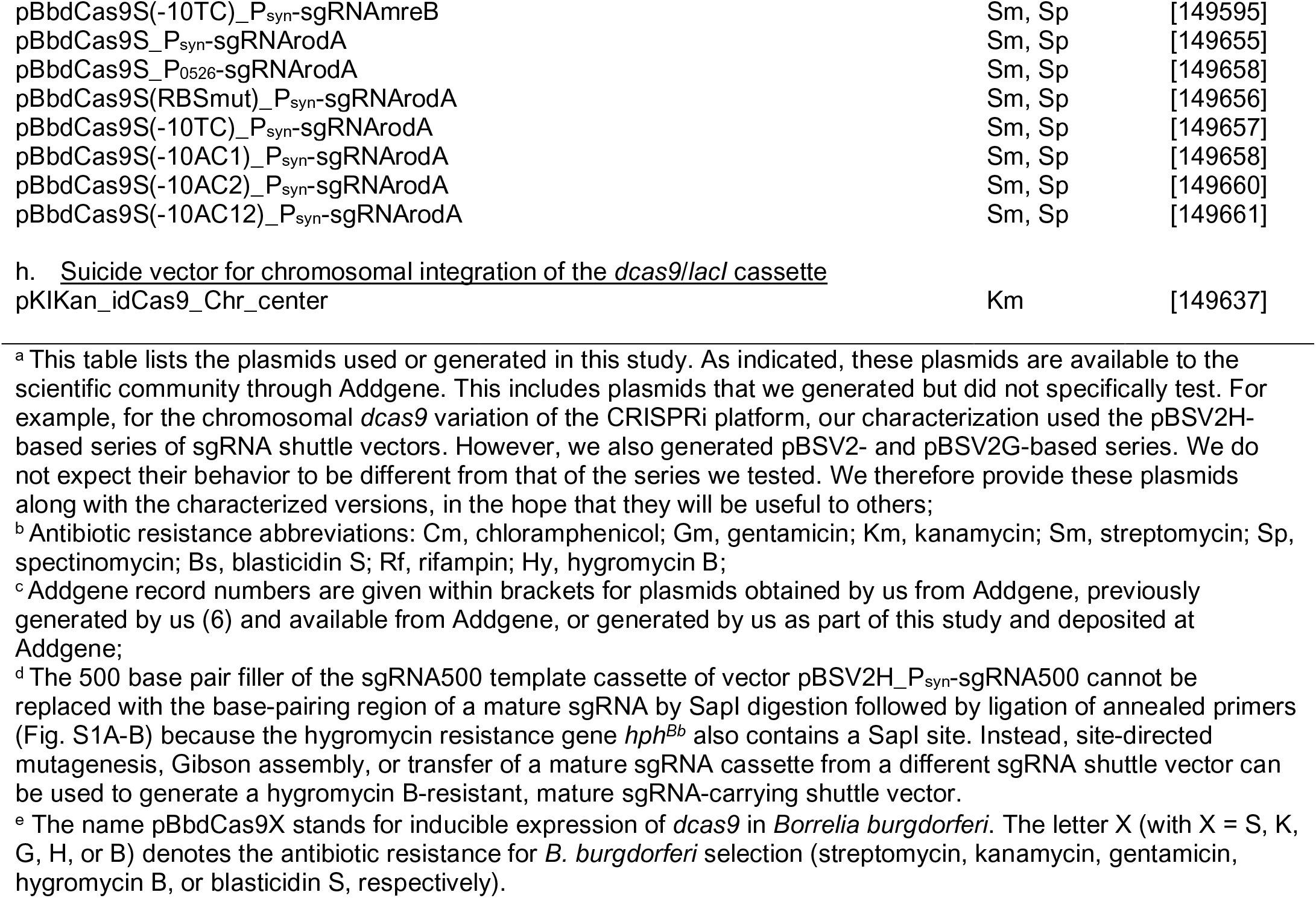
Plasmids used or generated in this study.

To express *dcas9*, we generated a *dcas9*/*lacI* cassette (Fig. 1B), which contains the *dcas9* gene controlled by the IPTG-inducible P*_pQE30_* promoter and a constitutively expressed *lacI* gene, both derived from plasmid pJSB252 (32). When the *dcas9*/*lacI* cassette is expressed from a *B. burgdorferi* shuttle vector, we hereafter refer to such a vector as a *dcas9* shuttle vector (Fig. 1D and Table 1). To facilitate cloning of a sgRNA cassette into the *dcas9* shuttle vector, we generated cloning intermediates that contain a rifampin resistance cassette placeholder (Fig. 1E and S1E and Table 1). Replacement of the rifampin cassette with sgRNA cassettes yielded all-in-one CRISPRi shuttle vectors (Fig. 1F and S1E and Table 1). To facilitate use of the CRISPRi platform in a variety of *B. burgdorferi* strains, including ones that already contain antibiotic resistance markers from prior genetic modifications, we generated five variations of the *dcas9* shuttle vectors, each carrying a different antibiotic resistance marker (Table 1).

We also inserted the *dcas9/lacI* cassette into an intergenic region of the chromosome of the infectious, transformable, B31-derived *B. burgdorferi* strain B31-A3-68 Δ*bbe02::P_flaB_-aadA* (33), generating strain CJW_Bb362 (Fig. 1G and Table 2). This strain allows for stable maintenance of the *dcas9/lacI* cassette in the absence of antibiotic selection. It requires transformation with a sgRNA shuttle vector (Fig. 1C) to generate a CRISPRi strain for protein depletion, while transformation with an empty shuttle vector yields a control strain.

**Table 2.**
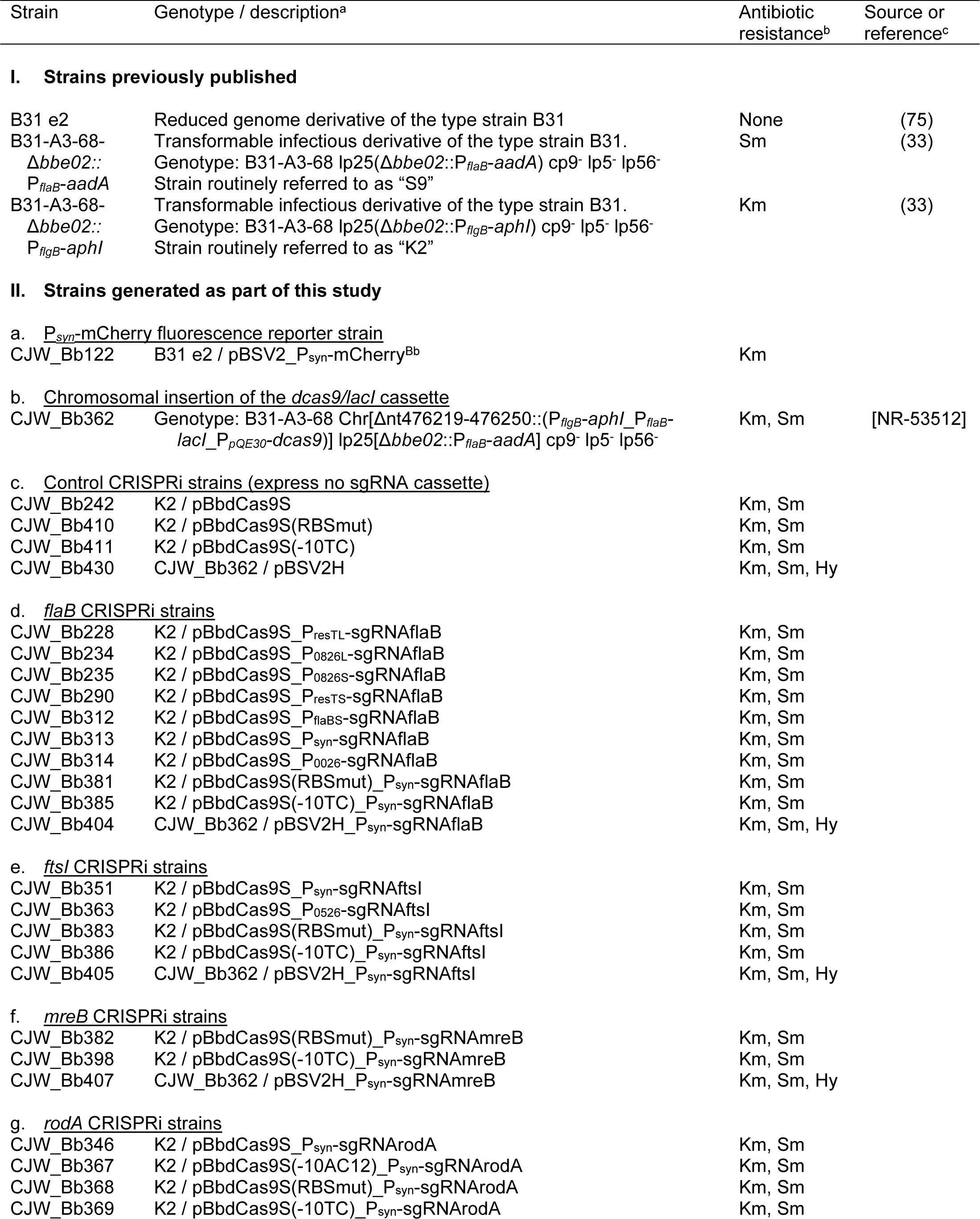
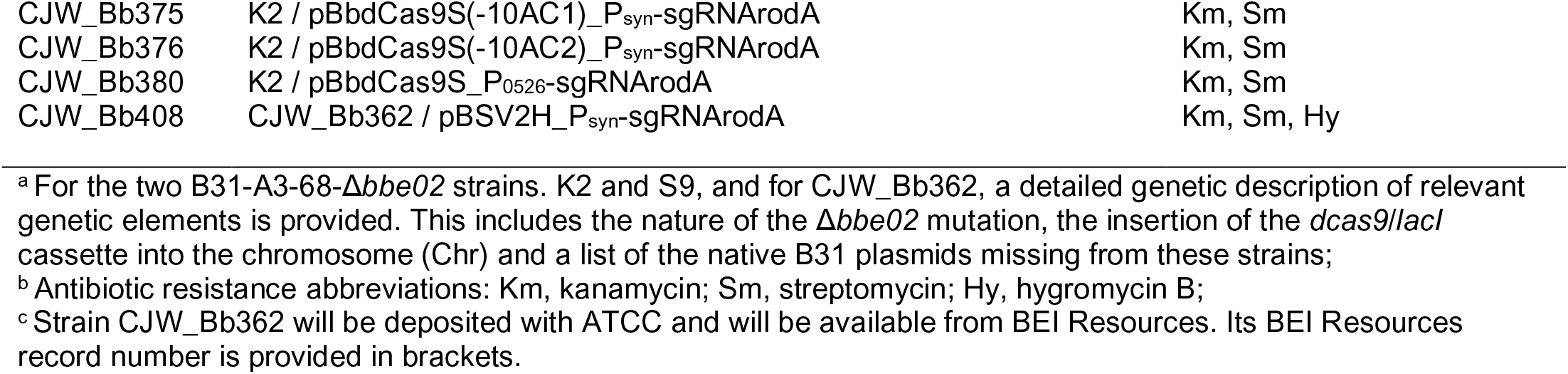
*B. burgdorferi* strains used or generated in this study.

Regardless of the CRISPRi version used (all-in-one CRISPRi shuttle vector or strain CJW_Bb362 transformed with an sgRNA shuttle vector), in the absence of IPTG (uninduced condition), cells of *B. burgdorferi* CRISPRi strains had their *dcas9* expression repressed by the binding of LacI to the *lacO* sites within the P*_pQE30_* promoter (Fig. 1H). IPTG addition to the cultures (induced condition) releases LacI from P*_pQE30_*, derepresses transcription from this promoter, and leads to synthesis of dCas9. Nonetheless, some background *dcas9* expression occurred in the absence of IPTG. As discussed below, this proved problematic when targeting certain *B. burgdorferi* genes. To decrease this basal *dcas9* expression, we mutated either the −10 promoter region of P*_pQE30_* or the ribosome-binding site upstream of the *dcas9* translational start codon in the background of the *dcas9* or the all-in-one CRISPRi shuttle vectors (Fig. 1H and S1F and Table 1). One of the promoter mutations (−10TC), as well as the ribosome-binding site mutation (RBSmut) proved effective in reducing background expression of *dcas9* and were analyzed in greater detail (see below).

Altogether, we generated and characterized four versions of the *B. burgdorferi* CRISPRi system. One is based on chromosomal expression of *dcas9* paired with plasmid-based expression of the sgRNA. The others are all-in-one plasmid-based versions that express both *dcas9* and the sgRNA from the same plasmid. The plasmid-based versions differ in whether the promoter that controls *dcas9* expression carries no mutation, a promoter mutation (−10TC) or the ribosomal binding site mutation (RBSmut).

### Inducible expression of *dcas9* in *B. burgdorferi*

To characterize *dcas9* expression in each of the four versions of our *B. burgdorferi* CRISPRi platform, we generated four corresponding sgRNA-free control strains (Table 2). We first measured, by quantitative real-time polymerase-chain reaction (qRT-PCR), their basal *dcas9* expression in the absence of IPTG induction. As expected, all control strains (CJW_Bb242, CJW_Bb410, and CJW_Bb411) harboring *dcas9* on a shuttle vector had higher levels of basal *dcas9* expression than the control strain (CJW_Bb430) carrying a chromosomal *dcas9* (Fig. 2A). Expression of *lacI* was also about five-fold higher in these strains than in the chromosomal *dcas9* strain (Fig. 2B). These levels could reflect copy-number differences between the shuttle vector and the chromosome, previously reported to be in a ratio of about 5:1 (34–36). They could also reflect changes in DNA topology and genomic context that are known to affect gene expression in *B. burgdorferi* (36–38).

**Figure 2.**
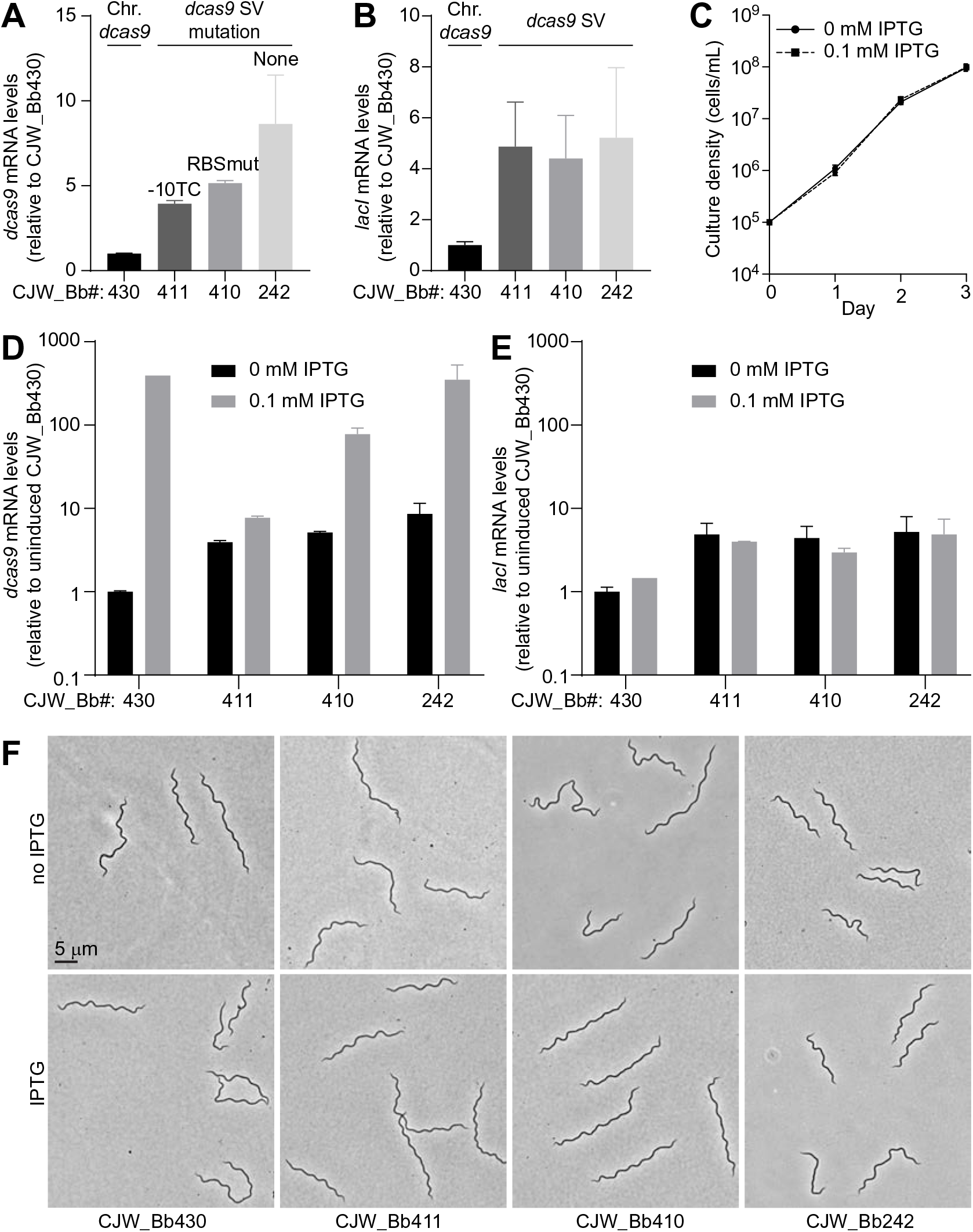
Characterization of *dcas9* expression in *B. burgdorferi*. **A.** Comparison of *dcas9* mRNA levels measured in the absence of IPTG in control strains that lack an sgRNA. Strain numbers are shown at the bottom. Relevant strain characteristics are marked on the graph. mRNA levels (measured by qRT-PCR) are shown relative to those in strain CJW_Bb430. Chr., chromosomal; SV, shuttle vector. **B.** Comparison of *lacI* mRNA levels measured in the absence of IPTG in the same samples as in panel A. **C.** Growth curve of strain CJW_Bb242 in the presence of 0 or 0.1 mM IPTG. Cell densities of three replicate cultures were counted daily. Shown are means ± standard deviations. **D.** IPTG-mediated induction of *dcas9* expression measured by qRT-PCR in the strains indicated below the graph. All values are reported relative to the levels in the uninduced strain CJW_Bb430. Measurements were obtained after one day of induction. **E.** *lacI* mRNA levels measured in the same samples as in panel D. A, B, D, and E. Shown are means ± standard deviations measured from two cultures. A single sample was measured for the induced CJW_Bb430 condition. **F.** Phase contrast images of cells of the indicated strains after two days of growth in the presence or absence of IPTG.

Basal levels of *dcas9* expression varied among the control strains carrying the *dcas9/lacI* cassette on a shuttle vector, with the highest level found in strain CJW_Bb242 (Fig. 2A), which has *dcas9* expression controlled by the parental P*_pQE30_* promoter (Fig. 1H). Mutation of the −10 region of this promoter yielded the largest drop in basal *dcas9* expression (Fig. 2A), likely reflecting lower promoter activity. The RBS mutation found in strain CJW_Bb410 also decreased basal *dcas9* expression levels (Fig. 2A), presumably reflecting lower stability of the *dcas9* mRNA due to lower ribosome occupancy. Translating ribosomes are known to protect bacterial mRNAs from degradation (39). The similar levels of *lacI* expression among the three strains that carry the *dcas9/lacI* cassette on a shuttle vector (Fig. 2B) suggest that the copy number of the shuttle vector, and thus that of *dcas9*, did not vary among these strains. Therefore, the mutations in the promoter and RBS decreased basal *dcas9* expression levels as intended.

Next, we determined appropriate induction conditions using strain CJW_Bb242 given it displayed the highest basal expression of *dcas9* (Fig. 2A). In this strain, induction of *dcas9* expression with 1 mM IPTG, a concentration known to elicit maximal expression from P*_pQE30_* in *B. burgdorferi* (32), resulted in slower growth in liquid culture and in semisolid BSK-agarose plates when compared to the no-IPTG condition (data not shown). This growth defect is not observed during the induction of expression of other genes (32). It is therefore likely due to overproduction of dCas9 and subsequent toxic effects associated with excessive nonspecific DNA binding, as observed in other bacteria (28, 40). Importantly, lowering the concentration of IPTG to 0.1 mM resulted in no discernable growth defect (Fig. 2C). We therefore used 0.1 mM IPTG to induce *dcas9* expression in all subsequent experiments.

We then measured the magnitude of induction of *dcas9* expression by IPTG. The highest magnitude, ∼400-fold, was observed in strain CJW_Bb430, which carries a chromosomal copy of the *dcas9/lacI* cassette (Fig. 2D). Among the shuttle vector-encoded *dcas9* versions, we observed the highest induction of ∼40-fold in the case of the unmutated promoter (Fig. 2D, strain CJW_Bb242). The RBSmut version displayed a ∼15-fold induction, while the −10 promoter mutation allowed for only a two-fold induction (Fig. 2D). The genetically-linked *lacI* gene experienced little change in its expression level in response to IPTG induction (Fig. 2E), as expected.

Finally, we imaged the strains after two days of exposure to IPTG and saw no notable changes in cell morphology between induced and uninduced cultures (Fig. 2F), further supporting the notion that these levels of *dcas9* induction are not toxic to the cells.

### *B. burgdorferi* genes targeted by CRISPRi

To test our CRISPRi platform, we targeted four *B. burgdorferi* genes: *flaB, ftsI, rodA,* and *mreB*. Depletion of their protein products is expected to yield morphological phenotypes, which are easily observable by microscopy. The *flaB* gene, which encodes flagellin, the major structural component of periplasmic flagella, is required for motility and for generating the characteristic flat wave morphology of *B. burgdorferi* cells (41, 42). FtsI is a transpeptidase that is required for septal peptidoglycan synthesis during cell division; its inhibition causes cell filamentation in *E. coli* (43–46). RodA, encoded by the *rodA* (*mrdB*) gene, is a transglycosylase active during lateral peptidoglycan synthesis in many rod-shaped bacteria (47). This lateral wall elongation is orchestrated by MreB, a bacterial actin homolog (48). When rod-shaped bacteria such as *E. coli* or *C. crescentus* lose RodA function, they grow into rounder shapes (49–51), as they do following MreB inactivation or depletion (52, 53).

Across them, the *flaB*, *rodA*, *mreB* and *ftsI* genes span two orders of magnitude of native expression levels (54), a range that covers a large subset of *B. burgdorferi* genes expressed during exponential growth in vitro (Fig. S2A). Furthermore, these genes are either the last gene in an operon or encode a monocistronic mRNA (Fig. S2B), rendering polar effects of CRISPRi unlikely. We designed sgRNAs targeting these four genes and cloned them either in sgRNA shuttle vectors or in all-in-one CRISPRi shuttle vectors (Table 1). These sgRNAs recognize sequences found in either the 5′ UTR or the protein-coding region of the targeted gene, near the translational start site (Fig. S2B). We introduced the sgRNA-expressing shuttle vectors into appropriate *B. burgdorferi* host strains, generating CRISPRi strains for depletion of FlaB, FtsI, MreB, and RodA (Table 2).

### Basal and induced CRISPRi activity in *B. burgdorferi*

In the absence of IPTG, clone generation, growth, and cell morphology were normal for the CRISPRi strains targeting *flaB,* regardless of the version of the CRISPRi platform used (Table S2). In contrast, upon transforming *B. burgdorferi* B31-A3-68 Δ*bbe02::P_flgB_-aphI* (also known as “K2”) (33) (Table 2) with all-in-one CRISPRi shuttle vectors containing an unmutated P*_pQE30_* promoter and targeting *rodA*, *mreB*, or *ftsI*, we observed phenotypes consistent with significant basal CRISPRi activity (Table S2). For example, we were unable to generate clones when transforming *B. burgdorferi* K2 with shuttle vectors targeting *mreB* (Table S2). We also observed delays in appearance of colonies in BSK-agarose plates when using shuttle vectors targeting *rodA* (Table S2). Even in the absence of IPTG, cells of the RodA depletion strain (CJW_Bb346) sometimes had normal flat wave morphologies but often displayed bulging (Fig. S3), consistent with a RodA depletion phenotype. We believe that the high basal plasmid-based expression of *dcas9* from the unmutated P*_pQE30_* promoter (Fig. 2A), combined with the constitutive expression of the sgRNA, led to formation of enough dCas9-sgRNA complexes to repress transcription of the targeted genes even in the absence of IPTG. When targeting *rodA*, introduction of the −10AC1, −10AC2, or −10AC12 mutations (Fig. S1F) into the P*_pQE30_* promoter in the context of the all-in-one CRISPRi shuttle vector did not fully abrogate the CRISPRi phenotype in the absence of IPTG induction (Table S2). We did not further analyze the strains carrying these constructs. In contrast, the −10TC or RBSmut versions of the all-in-one CRISPRi shuttle vector (Fig. 1H and Table 2), as well as the chromosomal *dcas9* version of the CRISPRi platform, allowed generation of strains that displayed no or weak phenotypic evidence of basal CRISPRi activity (Table S2).

We next characterized CRISPRi efficiency by qRT-PCR. In the absence of an sgRNA, the mRNA levels of *flaB, ftsI, mreB,* or *rodA* were not affected by IPTG addition (Fig. 3 and S4, gray bars). In the absence of IPTG, *flaB* mRNA levels were only slightly, if at all, reduced in strains constitutively expressing sgRNAflaB compared to the corresponding control strains lacking sgRNAflaB (Fig. 3A and S4A). In contrast, sgRNAs binding to *ftsI, mreB,* or *rodA* decreased their targets’ mRNA levels by ∼40% to ∼60% of those in the control strains, even without IPTG induction of *dcas9* expression (Fig. 3B-D and S4B-D). These lower mRNA levels appeared to be relatively well tolerated by the cells, as fewer than ∼1% of the cells in each population displayed morphological defects based on visual inspection (Table S2).

**Figure 3.**
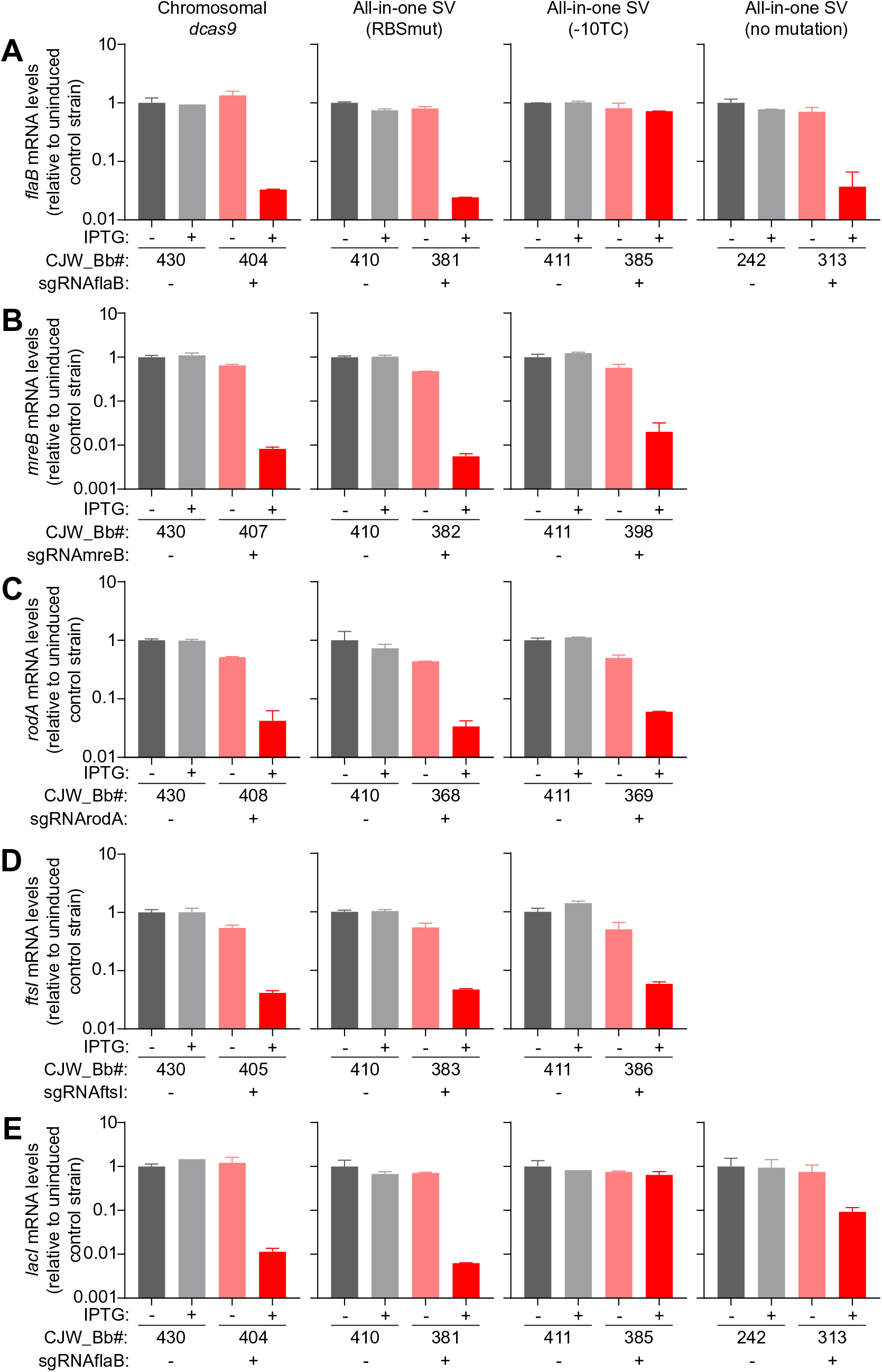
Effect of CRISPRi on targeted gene mRNA levels. **A.** *flaB*, **B.** *mreB,* **C.** *rodA*, **D.** *ftsI*, and **E.** *lacI* mRNA levels measured in the indicated control strains (gray) and CRISPRi depletion strains (pink and red) after one day of growth with or without IPTG. Shown are the means ± standard deviations measured from two cultures. The version of the CRISPRi platform carried by each set of strains is indicated above the corresponding column of graphs. SV, shuttle vector.

IPTG induction of *dcas9* expression for 24 hours in three of the four strains carrying sgRNAflaB (CJW_Bb404, CJW_Bb381 and CJW_Bb313) depleted *flaB* mRNAs by at least 95% of the levels measured in the corresponding control strains in the absence of IPTG (Fig. 3A). The depletion was maintained over two days of IPTG exposure (Fig. S4A). Induction of *dcas9* expression in these strains also depleted *lacI* mRNAs (Fig. 3E and S4E). This was expected, as *lacI* expression is controlled by P*_flaB_*, which contains the 5′ UTR of the *flaB* gene (7) and therefore the CRISPR site targeted by our sgRNA. In the remaining strain carrying sgRNAflaB (CJW_Bb385), *flaB* and *lacI* mRNA levels decreased noticeably less after IPTG induction (Fig. 3A, 3E, S4A, and S4E). Presumably, the weak induction of *dcas9* expression from the mutated (−10TC) *P_pQE30_* promoter (Fig. 2D) is insufficient to cause repression of both the *flaB* and *lacI* genes located on the chromosome and multicopy plasmid, respectively.

We also quantified *ftsI, mreB*, or *rodA* mRNA levels in the corresponding depletion strains grown in the presence of IPTG and compared them to controls lacking an sgRNA (Fig. 3B-D and S4B-D). In all strains, regardless of the version of the CRISPRi system present, induction of *dcas9* expression considerably depleted the mRNA of the targeted genes. The magnitude of the depletion after one day of IPTG exposure ranged from ∼95% for *rodA* and *ftsI* to ∼99% for *mreB* (Fig. 3B-D). Such low mRNA levels were still observed after two days following IPTG addition to the cultures (Fig. S4B-D).

### Phenotypic characterization of CRISPRi-mediated flagellin depletion

Despite the very high level of expression of *B. burgdorferi* flagellin (Fig. S2A), CRISPRi-mediated depletion of this protein was able to elicit a partial loss-of-function phenotype. Flagella impart to *B. burgdorferi* cells their motility and flat-wave morphology (41, 42), which we readily observed in uninduced or induced cells of control strains carrying no sgRNA (Movies S1 and S2 and Fig. 4A and S5A-C), or in uninduced cells of strains carrying sgRNAflaB (Movie S3 and Fig. 4B and S5A-D). After two days of IPTG exposure, cells of strain CJW_Bb313, which express sgRNAflaB from P*_pQE30_* on the shuttle vector, displayed varying degrees of motility defects (Movies S4-S8). While most cells retained some flat-wave morphology and twitching ability, they appeared straightened compared to their control counterparts (Fig. 4B). For the cells that were still able to move, most often the movement was slowed (Movie S4), reduced to a twitching pattern (Movies S5 and S6), or localized at the cell pole region (Movie S6). Some cells displayed little to no evidence of flagellum-based motion and instead appeared to simply display Brownian motion or drift in the medium (Movies S7 and S8). Some of these cells were almost completely straight, except for the occasional kink at the division site (Movie S8).

**Figure 4.**
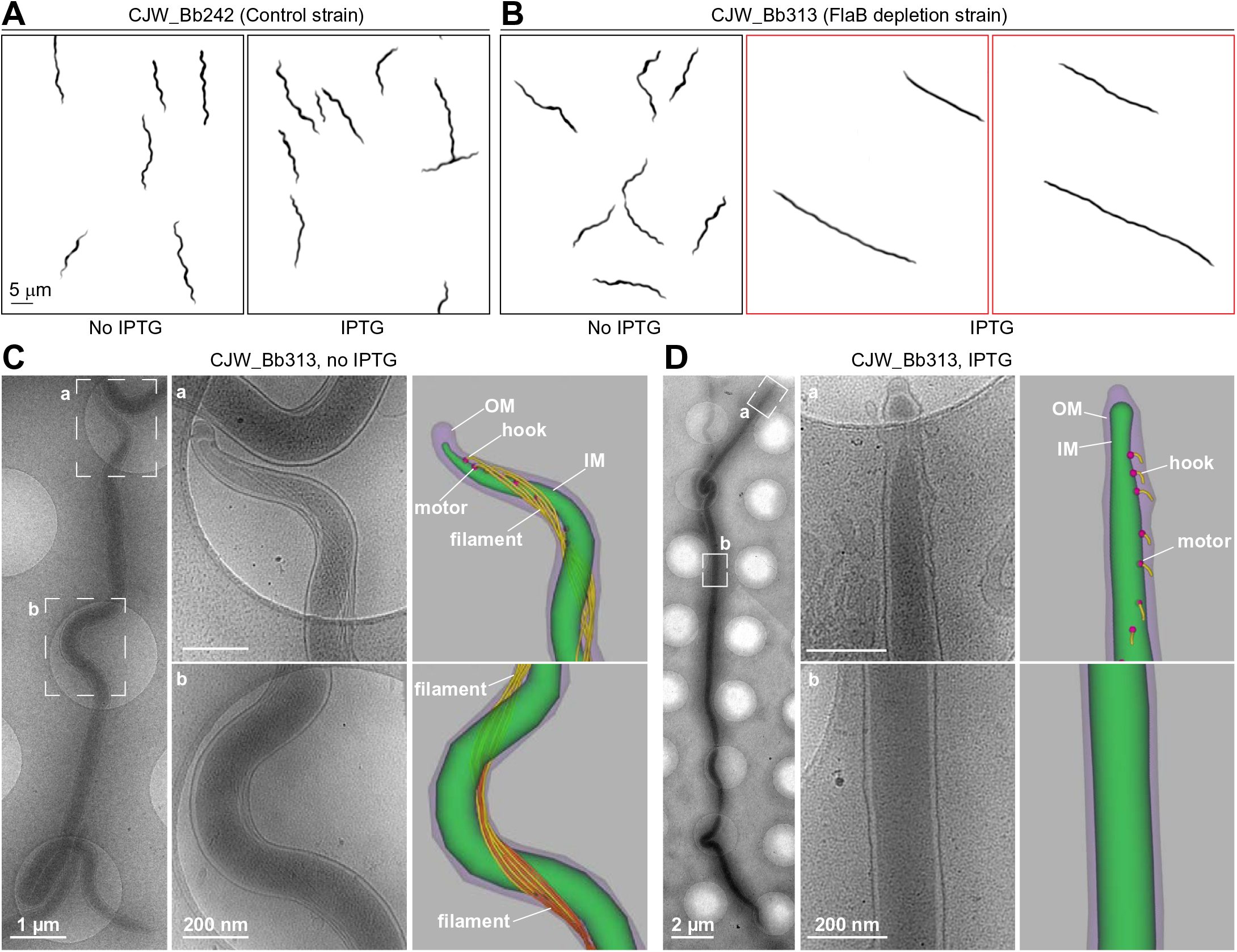
Phenotypic characterization of flagellin depletion. **A.** and **B.** Inverted darkfield images of strain CJW_Bb242 (A) or CJW_Bb313 (B) grown in the absence or presence of IPTG for two days. The flagellin depletion phenotype is highlighted by a red outline. **C.** Cryo-ET-based detection of periplasmic flagella in a cell of strain CJW_Bb313 grown in the absence of IPTG. Left: low magnification view of the entire cell. Center: high magnification views of the tip (a) and center (b) of the cell. Right: three-dimensional segmentation of the tip (panel a) and center (panel b) regions of the cell. In panel b, the flagella from one end of the cell are shown in yellow and the flagella from the other end are shown in orange. Also see Fig. S6A. **D.** Flagellin depletion assessed by cryo-ET in a cell of strain CJW_Bb313 after two days of IPTG exposure. Left: low magnification view of the entire cell. Center: high magnification views of the tip (a) and center (b) of the cell. Right: three-dimensional segmentation of the tip (panel a) and center (panel b) of the cell.

Expressing sgRNAflaB from seven different promoters of varied strengths resulted in a similar overall straightening of the cell body (Fig. 4B, S1D, and S5D and Tables 2 and S1). Our results therefore suggest that our CRISPRi system is robust to variation in sgRNA levels in *B. burgdorferi*. We also observed an overall straightening of the cell in IPTG-induced strains that carry either the chromosomal *dcas9* (CJW_Bb404, Fig. S5A) or the all-in-one CRISPRi shuttle vector with the RBS P*_pQE30_* mutation (CJW_Bb381, Fig. S5B). This is consistent with the significant *flaB* mRNA depletion observed in these strains (Fig. 3A and S4A). As expected, cells from strain CJW_Bb385, in which *flaB* transcript levels remained largely unaffected by IPTG induction (Fig. 3A and S4A), displayed normal morphology (Fig. S5C).

Flagellin depletion is expected to occur gradually over generations following IPTG induction, which could explain the mixed phenotypes we observed. In *B. burgdorferi*, multiple flagella are anchored near each cell pole (55, 56). They form bundles that extend from their subpolar anchors towards the center of the cell, where they overlap. We reasoned that retention of some wave-like cell shape in otherwise poorly motile or non-motile, flagellin-depleted cells may be caused by a decrease in the length or number of flagella. In this scenario, fewer flagella might exert enough force to bend the cell into a reduced-amplitude wave-like shape but not enough force to generate translational motion. To test these potential explanations, we imaged frozen-hydrated *B. burgdorferi* cells by cryo-electron tomography (cryo-ET). In the absence of IPTG, the cellular ultrastructure was indistinguishable from that previously reported (56, 57). Multiple flagella were attached near the cell poles (Fig. 4C, panel a, and Fig. S6A) and the flagellar bundles extended towards the middle of the cell, where they overlapped (Fig. 4C, panel b). In contrast, after exposure to IPTG for two days, the cells had no detectable flagellar filament around midcell (Fig. 4D and S6B, panels b), explaining the observed motility defects. At pole-proximal regions, flagellar hooks could be readily detected (Fig. 4D and S6A, panels a). However, no flagellar filament (Fig. 4D) or shorter filaments (Fig. S6A) could be detected at these subcellular regions, consistent with a strong depletion of FlaB.

### FtsI involvement in *B. burgdorferi* cell division

Knockdown of *ftsI* expression by CRISPRi for two days elicited significant cell filamentation in all the strains tested, in contrast to the uninduced cells of the same strains (Fig. 5A). Cell filamentation required expression of sgRNAftsI, as shown by cell length quantification (Fig. 5B). Cells almost 100 μm long, which is about five times the average length of a *B. burgdorferi* cell, were detected (Fig. 5B, bottom). We note that when immobilized between an agarose pad and a coverslip, as in our microscopy setup, longer cells are more likely to cross themselves or other cells, bend at a tight angle, or lie partly outside the field of imaging. In such cases, automated cell outline generation with the Oufti software package (58) was not possible and such cells were excluded from our measurements. Therefore, the cell length distributions measured in FtsI-depleted cultures likely underestimate the extent of cell filamentation present in the population. Regardless, our results implicate FtsI in *B. burgdorferi* cell division.

**Figure 5.**
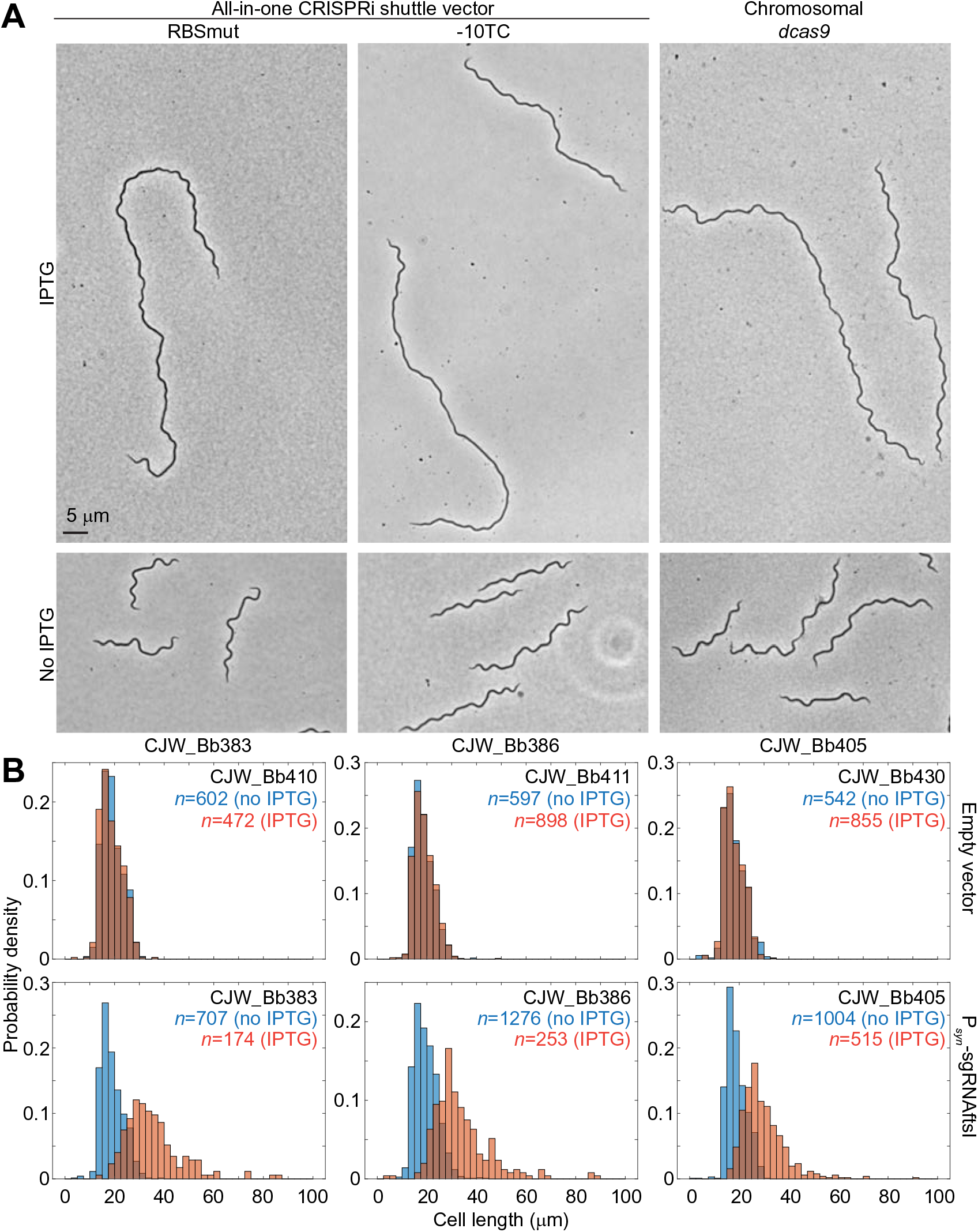
Phenotypic characterization of FtsI depletion. **A.** Phase contrast micrographs of cells from cultures of strains expressing sgRNAftsI after two days of *dcas9* induction with IPTG or in the absence of IPTG. **B.** Histograms depicting distributions of cell lengths measured in induced (0.1 mM IPTG for two days, orange) or uninduced (no IPTG, blue) cultures of the noted strains. The strains expressed either no sgRNA (top row) or sgRNAftsI (bottom row).

### Rod morphogenesis functions of MreB and RodA in *B. burgdorferi*

We imaged the MreB and RodA depletion strains before IPTG addition and daily after induction. We found that one day of MreB depletion was enough for significant cell bulging to develop (Fig. 6A). Bulging occurred predominantly at midcell (Fig. 6A, white arrowheads). In some cells, especially long ones, bulging was also apparent at ∼¼ and ∼¾ locations along the length of the cell (Fig. 6A, blue arrowheads). Our laboratory has previously shown that new peptidoglycan synthesis occurs at these subcellular locations in members of the *Borrelia* genus (59). The bulging phenotype at these locations therefore suggests that MreB is important for maintaining a constant cell width during insertion of peptidoglycan material at these specific sites (48). Furthermore, less pronounced cell body widening outside of these discrete locations, but encompassing longer segments of the cells, was also observed (Fig. 6A, yellow arrowheads). After two days of MreB depletion (Fig. 6A), the bulging phenotype became further exacerbated, with larger bulges at midcell and more pronounced laterally-spread cell body widening compared to the one-day time point. Overall, our findings establish a key role for MreB in *B. burgdorferi* cell morphogenesis.

**Figure 6.**
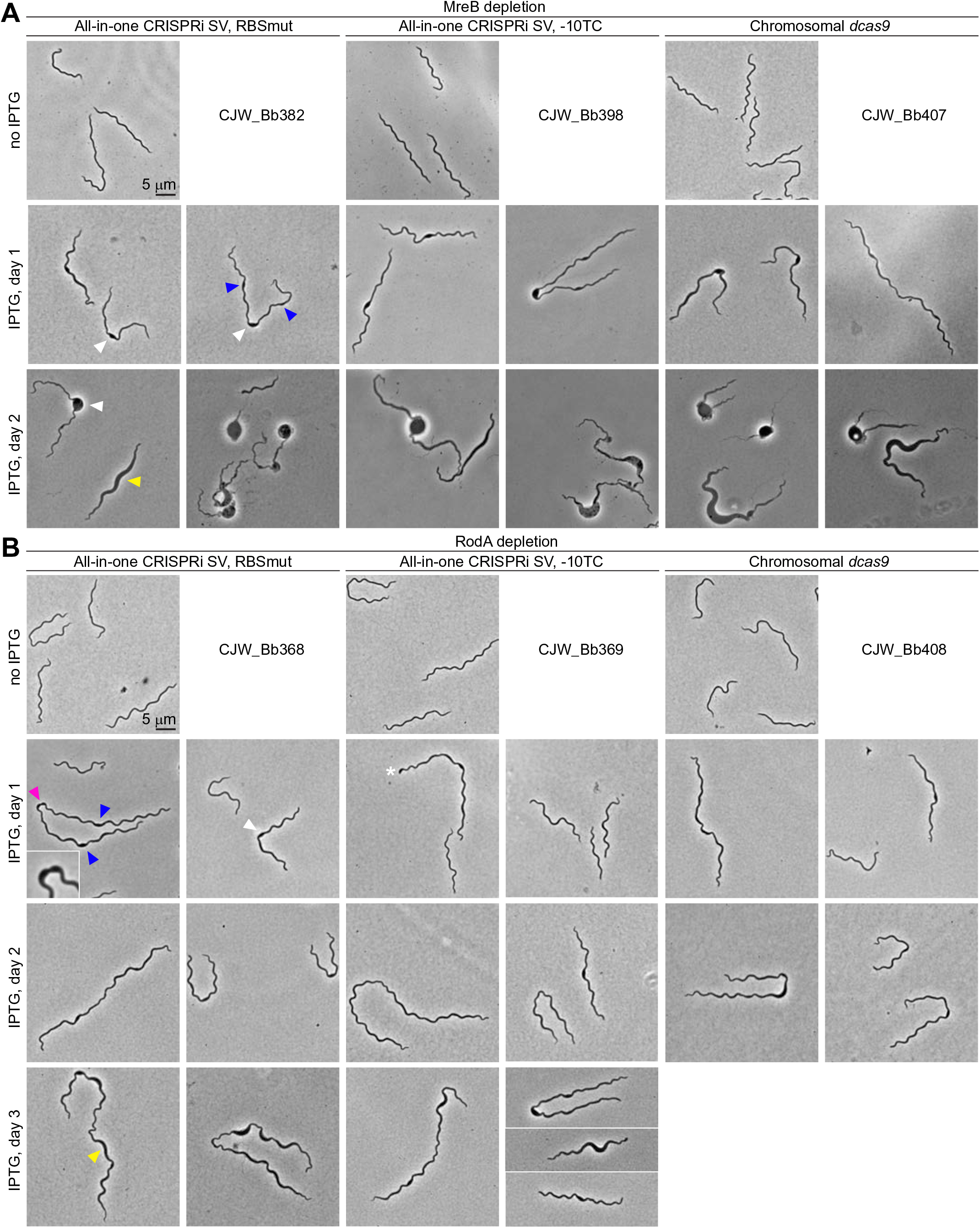
Phenotypic characterization of MreB and RodA depletion. **A.** and **B.** Phase contrast micrographs of cells from cultures of CRISPRi strains targeting *mreB* (**A**) or *rodA* (**B**). The marks show the following phenotypes: white arrowheads, cell bulging localized at midcell; blue arrowheads, cell bulging localized at approximately the ¼ and ¾ positions along the cell length; yellow arrowheads, cell widening extending along the cell length; pink arrowhead, a widened division site, shown in greater detail in the inset; and white asterisk, cell displaying an enlarged pole.

RodA depletion also elicited cell widening in *B. burgdorferi* (Fig. 6B). In the absence of IPTG, cells looked normal (Fig. 6B). Upon IPTG induction, bulging could be observed at midcell (Fig. 6B, white arrowhead) and, occasionally, at the ¼ and ¾ locations (Fig. 6B, blue arrowheads). Midcell bulging nevertheless permitted cell division to occur in some instances, as evidenced by the occurrence of deep constriction within a bulge (Fig. 6B, pink arrowhead and inset) and the occasional presence of rounded poles (Fig. 6B, white asterisk).

## DISCUSSION

In this study, we adapted the CRISPRi system for use in *B. burgdorferi* and tested its capabilities by targeting the motility and/or cell morphogenesis genes *flaB*, *ftsI*, *mreB*, and *rodA*. These genes span a broad range of native expression levels (Fig. S2A) (54), suggesting that CRISPRi will have broad applicability in the study of *B. burgdorferi*.

We created several variations of the CRISPRi platform. The all-in-one CRISPRi shuttle vectors can be introduced into any transformable *B. burgdorferi* strain, allowing comparison of behaviors across multiple genetic backgrounds and facilitating the pairing of CRISPRi-based gene depletion with other genetic methods. For example, using our system, the effects of depleting one gene product can be easily compared among otherwise isogenic strains that are wild-type, mutated, or complemented at a second gene locus. We further facilitated such pairing by generating CRISPRi shuttle vectors carrying each of five compatible antibiotic resistance markers (Table 1).

Another variant of our CRISPRi platform relies on two elements, a *dcas9/lacI* cassette stably integrated into the chromosome of a transformable derivative of the type strain B31 and an sgRNA expressed from a shuttle vector. The copy number of chromosomally expressed *dcas9* is expected to co-vary with the copy number of the targeted locus, whether it is located on the chromosome or on an endogenous plasmid. Prior studies have documented a ∼1:1 copy number ratio between various endogenous plasmids and the chromosome (34, 35, 60).

A common characteristic of all versions of our CRISPRi platform is that they only require a single shuttle vector transformation step. *B. burgdorferi* B31-derived strains that lack restriction modification enzymes are transformed by shuttle vectors at a higher frequency, in the range 10^−4^ to 10^−5^ (33), than the ∼10^−7^ frequency obtained using suicide vectors (7, 9). Thus, our CRISPRi platform offers an efficient complementary tool to homologous recombination-based methods for interrogation of gene function in *B. burgdorferi*.

In characterizing the CRISPRi platform, we found that low basal *dcas9* expression in the absence of IPTG combined with sizeable induction of *dcas9* expression upon IPTG addition appear to be important for broad functionality of the approach. Such is the case for the RBSmut version of the all-in-one CRISPRi shuttle vector and for the chromosomally-encoded *dcas9.* These CRISPRi platform versions yielded depletion phenotypes for all the genes we targeted. We recommend these versions in future CRISPRi experiments aimed at downregulating the expression of other genes.

Our CRISPRi approach allowed us to provide genetic insight into cell morphogenesis in *B. burgdorferi*. To our knowledge, neither deletion mutants nor transposon insertion mutants have been reported for *B. burgdorferi ftsI, mreB,* or *rodA* (61), potentially because these genes are essential for viability. Our inability to obtain clones while targeting *mreB* using the CRISPRi platform version that had the highest basal expression of *dcas9* supports this notion for *mreB*. We also tried to use A22 and MP265, two known small-molecule inhibitors of MreB (62–66), to study the function of this cytoskeletal protein in *B. burgdorferi*. Our attempts, however, proved unsuccessful, as *B. burgdorferi* appears to be resistant to chemical inhibition of MreB (see supplemental text in the combined supplemental material file and Fig. S7). Other examples of inhibitors widely used in other bacteria but not effective in *B. burgdorferi* include the transcription inhibitor rifampin, which acts on RpoB (67), and the peptidoglycan precursor synthesis inhibitor fosfomycin, which acts on MurA (43, 68). This list of important cellular functions that are apparently refractive to chemical inhibition in *B. burgdorferi* further underscores the utility of our easy-to-use and rapid CRISPRi genetic approach.

When we depleted FtsI, cells elongated into filaments, consistent with a conserved involvement of this protein in cell division (45, 69, 70). Depletion of RodA and MreB led to cell widening, reminiscent of loss-of-function phenotypes observed in other bacteria (50, 51, 53, 62, 70, 71). These results fit with the current model in which MreB orients lateral cell wall synthesis in such a way that a constant cell width is generated and propagated during growth (48). RodA, in turn, is an elongation specific transglycosylase (47) that contributes to the wall biosynthetic function organized by MreB (43, 48). Thus, the cell widening and bulging associated with MreB and RodA depletion in *B. burgdorferi* indicates that rod morphogenesis is controlled in this spirochete, at least in part, by some of the same actors as in other rod-shaped bacteria.

Bulging secondary to MreB depletion overwhelmingly occurred at sites of new peptidoglycan synthesis (59), primarily the midcell, but also the ¼ and ¾ positions along the length of the cells. The pattern of new peptidoglycan insertion in *Borrelia* species is peculiar compared to that seen in other bacteria, including other spirochetes such as those belonging to the *Treponema* and *Leptospira* genera (59). Those spirochetes elongate by inserting new peptidoglycan along the entire length of their cells. We do not yet know what mechanisms control the *Borrelia* pattern of new wall insertion. However, our observation that morphologic defects secondary to MreB depletion matched sites of new peptidoglycan insertion suggests a connection between MreB function and *B. burgdorferi*’s unusual pattern of cell wall growth.

Our results highlight the usefulness of a CRISPRi genetic approach. While this approach does not replace traditional homologous recombination-based methods, it offers some advantages as a complementary approach due to its procedural ease, speed, efficiency, and scalability. CRISPRi-based methods have proven to be particularly useful for the phenotypic characterization of genes essential for viability, for the simultaneous repression of multiple genes, and for high-throughput genome-wide studies (18, 19, 24, 28).

## MATERIALS AND METHODS

### Bacterial growth conditions

All *E. coli* strains were grown on LB agar plates or in Super Broth (35 g/liter bacto-tryptone, 20 g/liter yeast extract, 5 g/liter NaCl, 6 mM NaOH) liquid cultures at 30°C, with shaking. The following final concentrations of antibiotics were used for *E. coli* growth and selection: kanamycin at 50 μg/mL, gentamicin at 15-20 μg/mL, spectinomycin or streptomycin at 50 μg/mL, rifampin at 25 μg/mL for liquid selection or 50 μg/mL for growth on plates, and hygromycin B at 100-200 μg/mL. After heat shock or electroporation, *E. coli* transformants were allowed to recover in SOC medium (20 g/liter tryptone, 5 g/liter yeast extract, 10 mM NaCl, 2.5 mM KCl, 10 mM MgCl_2_, 10 mM MgSO_4_, 20 mM glucose) for 1 hour with shaking at 30°C, before plating. *E. coli* strain MC1000 (72) was grown overnight in LB liquid culture at 37°C with shaking, then diluted 1:1000 in BSK-II medium (see below), grown for another three hours, and finally treated with 50 μM MP265 or A22 (63) from a 1000X DMSO stock, or with DMSO alone, for one hour.

*B. burgdorferi* strains are listed in Table 2. They were grown at 34°C in a humidified incubator under 5% CO_2_ atmosphere (11, 13, 14). Liquid cultures were grown in Barbour-Stoenner-Kelly (BSK)-II medium (11), containing 50 g/liter bovine serum albumin (Millipore, #810036), 9.7 g/liter CMRL-1066 (US Biological, #C5900-01), 5 g/liter Neopeptone (Difco #211681), 2 g/liter Yeastolate (Difco #255772), 6 g/liter HEPES (Millipore #391338), 5 g/liter glucose (Sigma-Aldrich #G7021), 2.2 g/liter sodium bicarbonate (Sigma-Aldrich #S5761), 0.8 g/liter sodium pyruvate (Sigma-Aldrich #P5280), 0.7 g/liter sodium citrate (Fisher Scientific #BP327), 0.4 g/liter *N*-acetylglucosamine (Sigma-Aldrich #A3286), pH 7.60, and 60 mL/liter heat inactivated rabbit serum (Gibco #16120). BSK-1.5 medium contained 69.4 g/liter bovine serum albumin, 12.7 g/liter CMRL-1066, 6.9 g/liter Neopeptone, 3.5 g/liter Yeastolate, 8.3 g/liter HEPES, 6.9 g/liter glucose, 6.4 g/liter sodium bicarbonate, 1.1 g/liter sodium pyruvate, 1.0 g/liter sodium citrate, 0.6 g/liter *N*-acetylglucosamine, pH 7.50, and 40 mL/liter heat inactivated rabbit serum. Antibiotics were used at the following final concentrations for both liquid cultures and plates: kanamycin at 200 μg/mL, gentamicin at 40 μg/mL, streptomycin at 100 μg/mL, and hygromycin B at 300 μg/mL, respectively (6–8, 73). IPTG (American Bioanalytical #AB00841) was used at 0.1 mM final concentration from a stock of 1 M in water. Unless otherwise specified, cultures were always maintained in exponential growth, with culture densities kept below ∼5 x 10^7^ cells/mL. Cell density was determined by direct counting under darkfield illumination of samples placed in disposable hemocytometers, as previously described (6).

### *B. burgdorferi* transformation and clone isolation

Competent cells were generated as previously described (9). *B. burgdorferi* cultures containing exponentially growing cells were allowed to reach densities between 2 x 10^7^ cells/mL and ∼1 x 10^8^ cells/mL. Next, the cultures were centrifuged for 10 min at 10,000 x g and 4°C in 50 mL conical tubes, and the medium was removed by aspiration without disturbing the cell pellets. Cell pellets from 2-3 tubes were combined by resuspension in 40 mL cold EPS solution containing 93.1 g/liter sucrose (American Bioanalytical #AB01900) and 150 mL/liter glycerol (American Bioanalytical #AB00751) and centrifuged again. A second wash in 1 mL cold EPS was then performed, after which the cells were pelleted and resuspended in 50 to 100 μL of cold EPS for each 100 mL of initial culture. The competent cells were placed on ice or frozen at −80°C until use.

For cell transformation with a shuttle vector (4, 74), 25 to 50 μg of plasmid (in water) were mixed with 50-100 μL of competent cells of strain B31-A3-68-Δ*bbe02::*P*_flgB_*-*aphI* “K2” (33), B31 e2 (75), or CJW_Bb362, and electroporated (2.5 kV, 25 μF, 200 Ω, 2 mm gap cuvette). Electroporated cells were immediately recovered in 6 mL BSK-II and incubated overnight at 34°C. To insert the *dcas9/lacI* cassette into the chromosome, the pKIKan_idCas9_Chr_center plasmid (75 μg) was linearized in a 500 μL reaction volume containing 100 units of ApaLI enzyme for 4-6 hours at 37°C in CutSmart buffer (New England Biolabs). The DNA was then ethanol precipitated as previously described (76). The DNA pellet was dried in a biosafety cabinet, then resuspended in 25 μL water, chilled on ice, mixed with 100 μL of competent cells of the B31-A3-68-Δ*bbe02::*P*_flaB_-aadA* “S9” strain (33), and electroporated. The cells were allowed to recover overnight in 6 mL of BSK-II medium.

After overnight recovery, electroporated *B. burgdorferi* cells were plated in semisolid BSK-agarose medium. Three volumes of BSK-1.5 medium containing appropriate concentrations of antibiotics were equilibrated at 34-37°C, then briefly brought up to 55°C in a water bath and mixed with two volumes of 1.7% (w/v) agarose solution in water, also pre-equilibrated at 55°C. The mixed BSK-antibiotics-agarose solution (25 mL/plate) was added to 10 cm petri dishes containing aliquots of the electroporated *B. burgdorferi* cells and gently swirled to mix. The plates were chilled to room temperature for ∼30 min in a biosafety cabinet, then transferred to a humidified 5% CO_2_ incubator kept at 34°C for one to three weeks until colonies became clearly visible by eye. Alternatively, selection was performed in liquid culture by mixing 1 mL of electroporated cells with 5 mL of BSK-II medium containing appropriate concentrations of selective antibiotics. Beginning on the fifth day of selection, the cultures were visually inspected for growth by darkfield microscopy. Clones were isolated from these non-clonal, selected populations by limiting dilution in 96-well plates, as previously described (6) or by semi-solid BSK-agarose plating, as described above. Agarose plugs containing single colonies were removed from the plates and expanded in 6 mL of BSK-II medium containing selective antibiotics. Insertion of the *dcas9/lacI* cassette at the desired chromosomal location in strain CJW_Bb362 was confirmed by polymerase-chain reaction (PCR) analysis of isolated total genomic DNA (DNeasy Blood & Tissue Kit, Qiagen), using the following primer pairs: NT424 and NT425 (amplifies across the insertion site), NT591 and NT592 (amplifies within the kanamycin resistance gene *aphI*), NT681 and NT682 (amplifies within *lacI*), and NT683 and NT684 (amplifies within *dcas9*), respectively (see Table 3 for primer sequences). Except for strain CJW_Bb122 (which is not derived from an infectious strain), the plasmid complement of each isolated clone was determined using primer pairs previously described (77) and DreamTaq Green DNA Polymerase (ThermoFisher Scientific). Each clone was confirmed to contain all B31-specific plasmids, except lp5, cp9, and lp56, which are also absent from the parental strains (33). The clones were invariantly maintained in exponential growth and frozen at passage one or two.

**Table 3.**
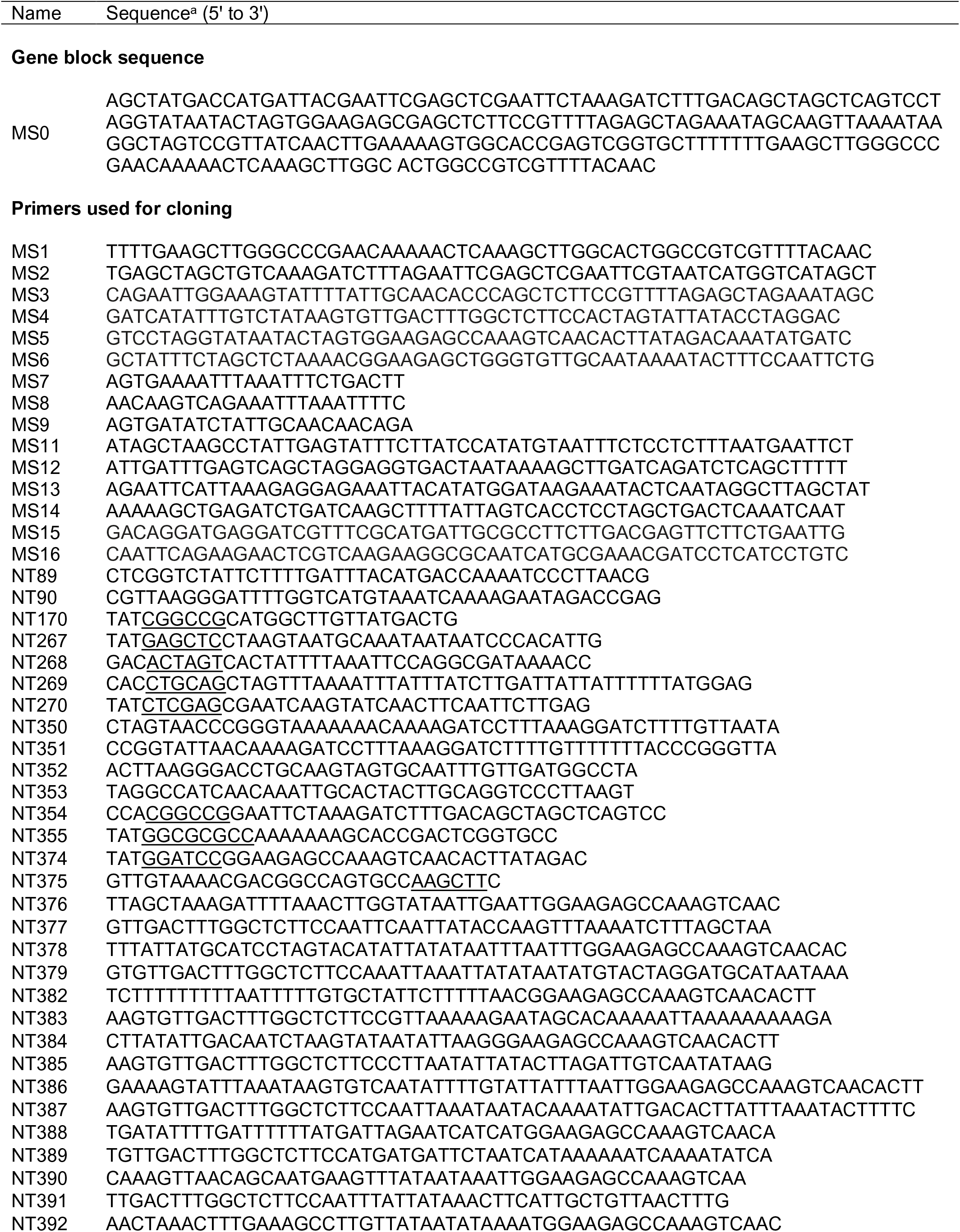
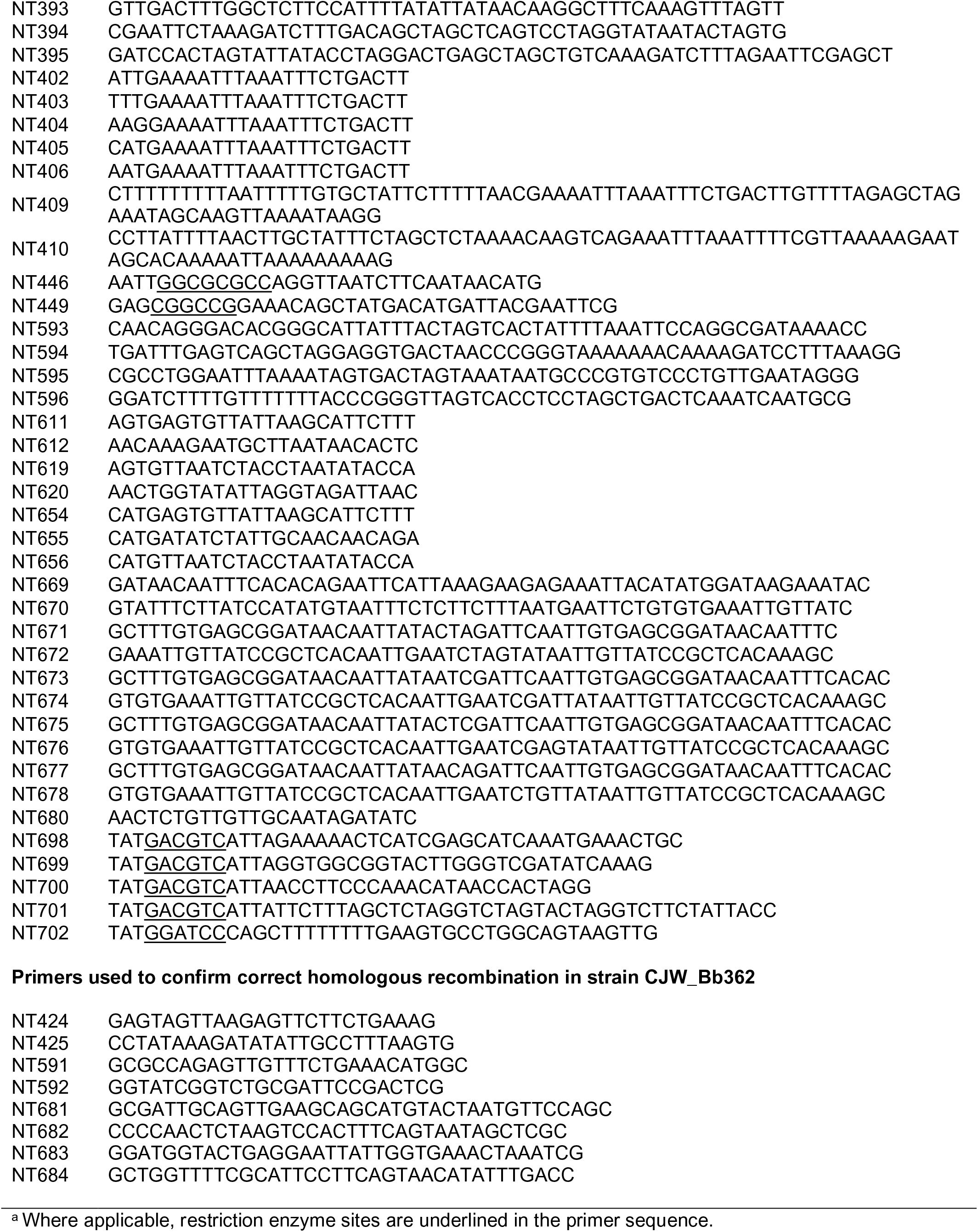
Synthetic DNA sequences used for cloning.

**Table 4.**
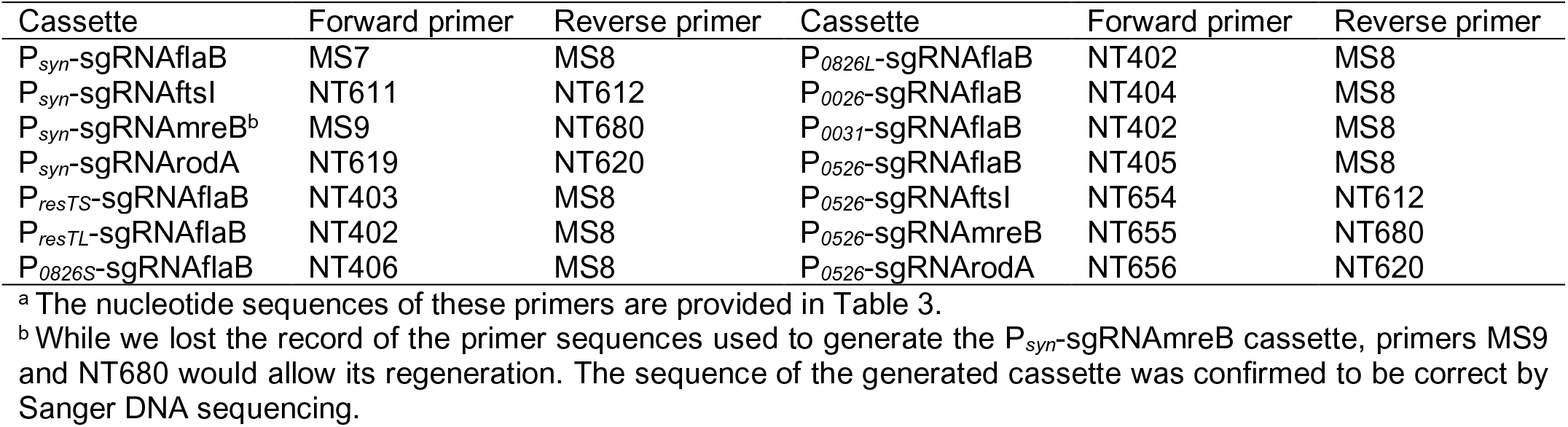
Primer pairs used to generate the mature sgRNA cassettes.

### DNA manipulations

#### General methods

Standard molecular biology techniques were used to generate the plasmids listed in Table 1. Oligonucleotide primers were purchased from Integrated DNA Technologies (IDT) and their sequences are provided in Table 3. PCR was performed using Platinum Hot Start PCR Master Mix or Phusion High-Fidelity DNA polymerase (ThermoFisher Scientific). Restriction endonucleases were obtained from New England Biolabs or ThermoFisher Scientific. Gel extraction was done using PureLink Quick Gel Extraction kit (ThermoFisher Scientific). Gibson Assembly Master Mix was procured from New England Biolabs. Agilent’s QuickChange Lightning Site-Directed Mutagenesis Kit was used according to the manufacturer’s protocol. New England Biolabs Electroligase or Quick Ligation Kits were used. Transformations of the cloning strains DH5α (Promega), NEB 5-alpha (New England Biolabs), or XL-10 Gold (Agilent) were done by electroporation or heat shock. Plasmids carrying the *dcas9/lacI* cassette were recovered and propagated in the NEB 5-alpha F’ *I^q^* strain (New England Biolabs). Cloning and/or propagation of plasmids containing the *dcas9/lacI* cassette in host *E. coli* strains that did not express *lacI^q^* often led to selection of clones that carried inactivating deletions within P*_pQE30_*, which abrogated *dcas9* expression in both *E. coli* and *B. burgdorferi*. This was likely due to high basal expression of *dcas9* in *E. coli* and associated non-specific toxic effects. Plasmid DNA was isolated using a Zyppy Plasmid Miniprep kit (Zymo Research) or Qiagen Plasmid Midi Kit. Correct DNA sequences of the relevant parts of the plasmids generated in this study were confirmed by Sanger sequencing at Quintarabio or at the Yale Keck Biotechnology Resource Laboratory.

#### Generation of sgRNA cassettes

The cloning template for an sgRNA cassette, modeled after a previous report (19), contains the following features: a promoter, a 500 bp filler sequence, a dCas9 handle region, and a transcriptional terminator sequence (Fig. 1A). To generate the P_syn_-sgRNA500 template cassette, a gene block (MS0, Table 3) containing the promoter P*_syn_*, the dCas9 handle region, and the transcriptional terminator, as well as homology regions to the *B. burgdorferi* shuttle vectors, was synthesized at Integrated DNA Technologies. This gene block was Gibson-assembled with the backbones of pBSV2 (78) and pBSV2G (8), which were obtained by PCR amplification using primers MS1 and MS2, respectively. The resulting plasmids were PCR-amplified using primers MS3 and MS4, and Gibson-assembled with a ∼500 bp filler DNA sequence. This filler was obtained by PCR amplification of the luciferase gene of pJSB252 (32) using primers MS5 and MS6.

To place sgRNA expression under the control of constitutive *B. burgdorferi* promoters (Table S1), the following steps were taken. The sgRNA500 sequence was PCR amplified with primers NT374 and NT375, digested with BamHI and HindIII, and cloned into the BamHI and HindIII sites of shuttle vectors generated previously (6) that contain the following promoters between their SacI and BamHI sites: P*_flaB_*, P*_resT_*, P*_0026_*, P*_0031_*, P*_0526_*, or P*_0826_*. Next, the 5′ UTR contained within these promoter sequences (Table S1), as well as the BamHI site, were deleted by site-directed mutagenesis. The following primer pairs were used on the appropriate template: NT382 and NT383 generated the P*_flaBS_*-sgRNA500 cassette; NT378 and NT379 generated P*_resTS_*-sgRNA500; NT376 and NT377 generated P*_resTL_*-sgRNA500; NT392 and NT393 generated P*_0826S_*-sgRNA500; NT390 and NT391 generated P*_0826L_*-sgRNA500; NT384 and NT385 generated P*_0026_*-sgRNA500; NT388 and NT389 generated P*_0526_*-sgRNA500, and NT386 and NT387 generated P*_0031_*-sgRNA500.

To generate mature sgRNA cassettes targeting specific genes, the 500-bp filler was excised from plasmids containing the template sgRNA500 cassette by digestion with SapI, BspQI, or LguI. Pairs of primers (Tables 3 and 4) were annealed, generating a short dsDNA sequence with overhangs complementary to the overhangs generated by SapI, BspQI, or LguI digestion of the plasmid. The digested plasmid and annealed primers were ligated. To obtain pBSV2G_P_flaBS_-sgRNAflaB, site-directed mutagenesis was performed on pBSV2G_P_flaBS_-sgRNA500 using primers NT409 and NT410.

#### Choice of sgRNA target base-pairing sequence

The coding sequence of a gene of interest, in the 5′ to 3′ orientation and including ∼100 bp upstream of the gene (to ensure that the 5′ UTR, if present, was included), was imported into the Geneious R10 software package. CRISPR sites were then identified using the “Find CRISPR sites” feature of the software. The required parameters were a 20-nucleotide base pairing region upstream of an NGG-3′ PAM site. CRISPR target sites were selected for further evaluation based on the following criteria: (i) they mapped within the target gene’s 5′ UTR or within its protein-coding region close to the translational start site and (ii) their orientation was opposite that of the coding region, thus ensuring that the sgRNA targets the non-template strand (18, 19). Next, the NCBI webtool blastn was used to compare the CRISPR target site sequence against the complete *B. burgdorferi* B31 genome sequence (79) to rule out off-target binding. Primer pairs were then designed to encode sequences complementary to the CRISPR target sites, minus the PAM. A guanine base was added 5′ to the base-pairing sequence to ensure similar efficiency of transcription of the various sgRNAs and to account for the purine preference at the +1 position of the TSS observed across several bacterial genomes (80, 81). Finally, the primers were designed to also generate, upon annealing, overhangs compatible with the SapI-digested plasmids containing template sgRNA cassettes.

#### Transfer of sgRNA cassettes

Among the shuttle vectors that do not contain the *dcas9/lacI* cassette, sgRNA cassettes were transferred as SacI/FspI restriction fragments. P*_syn_*-containing cassettes (e.g., P_syn_-sgRNAflaB) were inserted into pBbdCas9 vectors as AscI/EagI digests of PCR products generated using primers NT354 and NT355 and the appropriate template DNA. All other cassettes were inserted into pBbdCas9 vectors as AscI/EagI digests of PCR products generated using primers NT355 and NT449. The sgRNA cassettes were transferred among various pBbdCas9 vectors (see below) as either AscI/EagI or EagI/XmaI digests.

#### Updated sequence of pBSV2H

During this work, we discovered that pBSV2H, which we generated as part of our previous study (6), contains a duplication of its dual rifampin-hygromycin B resistance cassette. This was confirmed by quality control tests performed at Addgene. We have therefore updated the sequence of the construct on Addgene’s product page (#118229). All pBSV2H-derived plasmids generated in this study (Table 1) also contain this duplication. However, based on the normal behavior of the *B. burgdorferi* strains generated with these plasmids (see results obtained using chromosomal *dcas9* CRISPRi strains), we believe that the antibiotic cassette duplication does not affect the functionality of these vectors. Nevertheless, we generated a version of this shuttle vector that carries only one copy of the dual antibiotic resistance cassette, which we named pBSV2H_2 (Table 1).

#### Generation of all-in-one CRISPRi shuttle vectors

The luciferase gene of pJSB252 (32) was replaced with *dcas9* as follows. The pJSB252 backbone was PCR-amplified with primers MS11 and MS12. The *dcas9* gene encoding the catalytically inactive protein (18) was PCR-amplified from plasmid pdCas9-bacteria using primers MS13 and MS14. The two fragments were Gibson-assembled. Next, a silent mutation was introduced into the *lacI* gene to remove the SapI site. This was done by site-directed mutagenesis using NT352 and NT353, yielding plasmid pBbdCas9S. We note that the *dcas9* gene present in our constructs carries a mutation that results in a M1169L amino acid change, but *dcas9* rermains functional despite this change. The *arr2* rifampin-resistance gene was then PCR-amplified from pBSV2B (6) using primers NT170 and NT446, digested with AscI and EagI, and cloned into the AscI/EagI backbone of pBbdCas9S to yield pBbdCas9S_arr2. P*_flgB_*-driven antibiotic resistance marker cassettes were generated as follows: primers NT698 and NT702 were used to PCR amplify the kanamycin cassette of pBSV2_2; NT699 and NT702 were used to amplify the gentamicin cassette of pBSV2G_2; NT700 and NT702 were used to amplify the blasticidin S cassette of pBSV2B; and NT701 and NT702 were used to amplify the hygromycin B cassette of pBSV2H. The resulting PCR products were digested with BamHI and AatII and ligated into the backbone of pBbdCas9S_arr2 obtained following sequential BglII and AatII digestion. This process generated the following plasmids: pBbdCas9K_arr2 (kanamycin resistant), pBbdCas9G_arr2 (gentamicin resistant), pBbdCas9B_arr2 (blasticidin S resistant), and pBbdCas9H_arr2 (hygromycin B resistant) (Table 1).

#### Altered regulation of *dcas9* expression

The following primer pairs were used to modify, by site-directed mutagenesis, the DNA region upstream of the *dcas9* coding sequence. NT669 and NT670 were used to mutate the ribosomal binding site, generating RBSmut constructs (Fig. 1H). The rest of the mutagenesis reactions modified the −10 region of the *dcas9* promoter, P*_pQE30_*, as follows: NT677 and NT678 were used to generate the −10TC construct, NT671 and NT672 for the −10AC1 construct, NT673 and NT674 for the −10AC2 construct, and NT675 and NT676 for the −10AC12 construct (Fig. 1H and S1F).

#### Suicide vector for chromosome integration of the *dcas9/lacI* cassette

The following gene segments were assembled through a series of intermediate constructs. (i) The *aphI* gene of pCR2.1 TOPO was deleted by Gibson assembly using primers MS15 and MS16. The *bla* gene of the resulting backbone was deleted by site-directed mutagenesis using primers NT89 and NT90. The resulting backbone retains the *E. coli* origin of replication of pCR2.1 and its multicloning site. (ii) The antibiotic resistance cassette was assembled into this backbone by linking P*_flgB_* (flanked by SacII and NdeI sites), the *aphI* gene (flanked by NdeI and XmaI sites), and the *flaB* terminator, obtained by annealing primers NT350 and NT351 and inserting them between sites SpeI and XmaI. (iii) The sequence from nucleotide 474180 to nucleotide 476218 of the B31 chromosome was amplified with primers NT267 and NT268 and cloned as a SacI/SpeI fragment. (iv) The sequence from nucleotide 476251 to nucleotide 478279 of the B31 chromosome was amplified with primers NT269 and NT270 and cloned as a PstI/XhoI fragment. The backbone, antibiotic resistance cassette, and the two homology regions allow insertion by double cross-over into the B31 chromosome between nucleotide 476218 and nucleotide 476251 in the intergenic region between genes *bb0456* and *bb0457*. This suicide vector sequence was amplified with primers NT593 and NT594, while the *dcas9/lacI* cassette was PCR-amplified using primers NT595 and NT596 from pBbdCas9S_P_resTS_-sgRNAflaB. These two PCR products were Gibson-assembled to yield pKIKan_idCas9_Chr_center.

#### pBSV2_Psyn-mCherry^Bb^

Primers NT394 and NT395 were annealed and ligated into the SacI/BamHI backbone of pBSV2_P_resT_-mCherry^Bb^ (6).

### RNA isolation and qRT-PCR

Exponentially growing cultures of the CRISPRi strains were counted and diluted to 10^6^ cells/mL for next-day harvesting, or to 10^5^ cells/mL for day-two harvesting. For situations in which IPTG induction caused growth defects, the culture induced for two days was also started at a density of 10^6^ cells/mL to ensure that similar amounts of RNA were obtained as in the non-induced culture at the time of harvest. Dilutions were carried out in 25 mL of BSK-II medium with or without 0.1 mM IPTG. At 24 or 48 hours post-induction, bacteria were harvested by centrifugation of the 25 mL culture for 10 minutes at 4300 x g and room temperature in a swinging bucket centrifuge. Supernatant was aspirated and the pellet was resuspended in 400-500 μL HN buffer (50 mM NaCl, 10 mM HEPES, pH 8.0) (82). The suspension was then immediately added to 1 mL RNAprotect Bacteria Reagent (Qiagen), vortexed, incubated for 5 minutes at room temperature, and centrifuged at 10,000 x g and room temperature for 10 minutes. The supernatant was aspirated and the pellet was stored at − 80°C until RNA could be extracted using an enzymatic lysis and proteinase K digestion method (Protocol 4 in the RNAprotect Bacteria Reagent manual), followed by purification using the RNeasy Mini kit (Qiagen). A DNase digestion step was then performed using the TURBO DNA-free kit (ThermoFisher Scientific) and purified RNA was stored at −80°C.

The KAPA SYBR FAST One-Step qRT-PCR Master Mix Kit was used to quantify transcript levels. Reactions were done in duplicate or triplicate, using 10 ng total RNA per reaction. At least one additional reaction was performed for each sample without the addition of the reverse transcriptase mix. The Cq values obtained from these control reactions confirmed that none of our results could be explained by DNA contamination of the RNA sample. The following protocol was used on a Bio-Rad iCycler: reverse transcription (5 min at 42°C); enzyme activation (3 min at 95°C); 40 cycles of annealing, extension, and data acquisition (3 sec at 95°C, 20 sec at 60°C); and a melt curve analysis (55 to 95°C in 0.5°C increments). The primers used to PCR amplify the various targets are provided in Table 5. The amount of each transcript of interest was normalized to the level of *recA* transcript in the same sample, and then expressed relative to the level of target transcript in the control sample, as previously described (83).

**Table 5.**
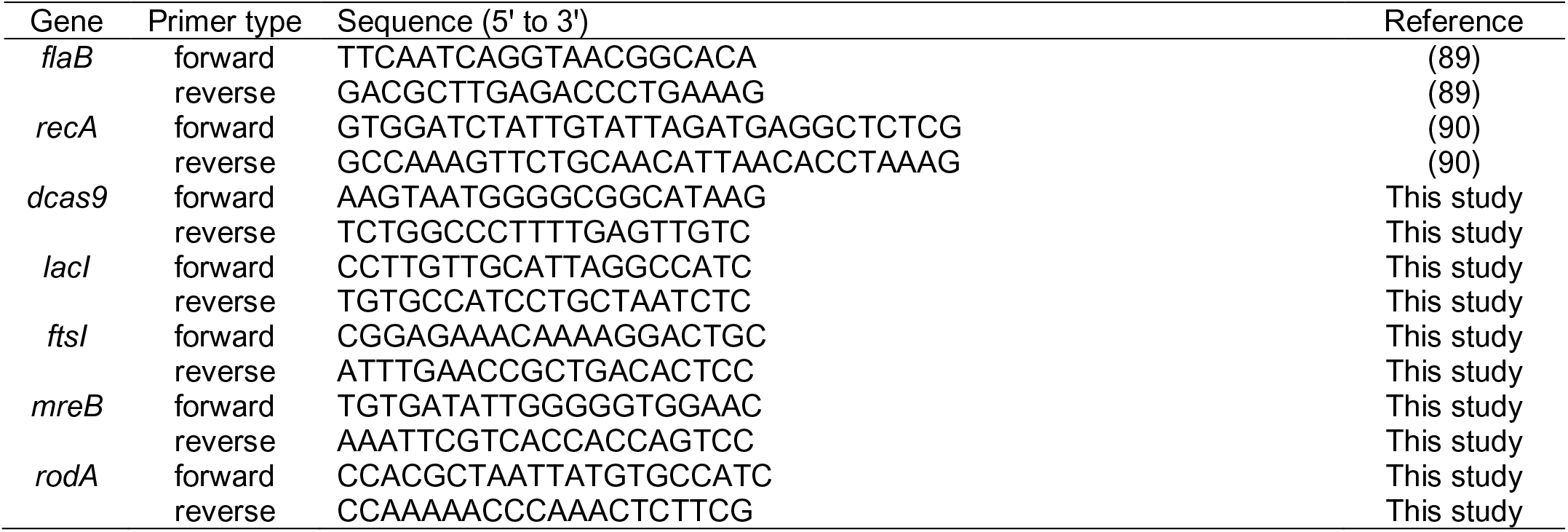
Primers used for qRT-PCR.

### Microscopy

Routine darkfield imaging of cultures was accomplished using a Nikon 40X 0.55 numerical aperture (NA) Ph2 phase-contrast air objective mounted on a Nikon Eclipse E600 microscope equipped with a darkfield condenser ring. Darkfield images and movies were acquired on a Nikon Eclipse Ti microscope equipped with a 40X 0.60 NA objective, a Nikon dry darkfield condenser (0.80-0.95 NA), and a Hamamatsu Orca-Flash4.0 V2 digital complementary metal-oxide semiconductor (CMOS) camera. The same microscope was used to obtain phase contrast images using a 100X Plan Apo 1.45 NA Ph3 phase-contrast oil immersion objective and a Ph3 condenser ring. The microscope was controlled using the Nikon Elements software. Fluorescence imaging of strain CJW_Bb122 was done on a Nikon Eclipse Ti microscope with the following features: a 100X Plan Apo 1.40 NA Ph3 phase-contrast oil objective, a Hamamatsu Orca-Flash4.0 V2 CMOS camera, a Sola light engine (Lumencor), an mCherry/TexasRed fluorescence filter cube containing an ET560/40x excitation filter, a T585lpxr dichroic mirror, and an ET630/75m emission filter (Chroma), and Metamorph software (Molecular Devices). Images were processed using Nikon Elements Software, Metamorph software, or Fiji software (84).

### Image analysis

Cell outlines were generated based on phase-contrast images using Oufti, our open-source analysis software package (58). The raw outlines were curated as follows: (i) outlines assigned to cells that crossed other cells, folded upon themselves, or were only partially present in the imaged frame, as well as outlines assigned to image components other than cells, were manually removed; (ii) outlines were manually extended to the tips of the cells where appropriate; (iii) outlines were manually added to cells whose geometry could support an outline but whose outlines were not generated automatically by Oufti; and (iv) when clear dips in the phase signal indicated an outer membrane bridge between two cytoplasmic cylinders that had separated as part of the cell division process, the two sides of the cell were treated as independent cellular units. In such cases, separate outlines generated by Oufti were left in place, while single outlines were manually split at the point at which the phase signal was observed to dip. Finally, all outlines were refined using the Refine ALL function of the Oufti software, and the fluorescence signal data was added to the outlines. Fluorescence quantification was done as previously described, using the MATLAB script addMeshtoCellList.m and the function CalculateFluorPerCell.m (6). Cell lengths were extracted using the MATLAB script get_um_lengths.m (85), and values below 1 μm were excluded from the analysis.

### Cryo-electron tomography and three-dimensional visualization

*B. burgdorferi* cultures growing exponentially in BSK-II medium to no more that 5×10^7^ cells/mL were pelleted for 10 minutes at 3,000 x g at room temperature. The pellet was gently resuspended in a small volume (50-100 μL) of BSK-H (Sigma-Aldrich, #B8291). The cell suspension was then mixed with 10-nm colloidal gold fiducial markers. Five microliters of cell suspension were deposited on holey carbon electron microscopy grids (200 mesh, R2/1, Quantifoil), which had been freshly glow-discharged for ∼30 seconds. The grids were then blotted with filter paper and rapidly frozen in liquid ethane using a homemade gravity-driven plunger apparatus. The frozen-hydrated specimens were transferred to a 300 kV Titan Krios electron microscope (ThermoFisher Scientific) equipped with a K2 Direct Electron Detector and energy filter (Gatan). SerialEM (86) was used to collect single-axis tilt series around −5 μm defocus, with a cumulative dose of ∼70 e^−^/Å covering angles from −51° to 51° with a 3° tilt step. Images were acquired at 26,000X magnification with an effective pixel size of 5.457 Å at the specimen level. All recorded images were first drift corrected using the MotionCor2 software (87) and then stacked by the software package IMOD (88). In total, 11 tilt series were aligned and reconstructed using IMOD. Three-dimensional models of the flagella and cells were manually segmented and visualized by IMOD.

### DATA AVAILABILITY AND ACCESSION NUMBERS

Plasmids generated in this study (and their sequences) are available through Addgene under the accession numbers listed in Table 1. Strain CJW_Bb362 will be made available through ATCC’s BEI Resources collection (Item # NR-53512). Reasonable requests for all other *B. burgdorferi* strains generated in this study shall be honored by the Jacobs-Wagner lab. The MATLAB code used to process cell fluorescence data can be downloaded from GitHub (85). MATLAB code, including dependencies, are provided at GitHub under https://github.com/JacobsWagnerLab/published.

## ACKNOWLEDGEMENTS

We thank Dr. Brandon Jutras for providing an early intermediate plasmid used in the generation of the pKIKan_idCas9_Chr_center plasmid, Nicholas Jannetty for assistance in generating pBSV2_P_syn_-mCherry^Bb^, and the members of the Jacobs-Wagner lab for critical reading of the manuscript. We also thank Dr. Jon Blevins (University of Arkansas for Medical Sciences) for sharing the pJSB252 plasmid and Dr. Will Arnold (Addgene) for sharing the quality control data on plasmid pBSV2H.

## SUPPLEMENTAL MATERIAL

### SUPPLEMENTAL TEXT

#### Pharmacologic attempts to study MreB function in *B. burgdorferi*

Specific small-molecule inhibitors can be valuable tools in biological investigations. For instance, the compounds A22 and MP265 inhibit the bacterial actin homolog, MreB, causing cell rounding of *E. coli* and other rod-shaped bacteria (1–5). A22 is active in the related spirochete *Leptospira biflexa*, where it causes cell bulging (6). However, the effects of MreB inhibitors on *B. burgdorferi* morphology have not been reported.

When we exposed *B. burgdorferi* strain K2 for two days (about seven generations) to MP265 at 50 μM, the typical dose of MreB inhibitor used in other bacteria (1, 2, 6), we observed no detectable morphological changes (Fig. S7A). Increasing the MP265 dose to 500 μM or treating *B. burgdorferi* with 500 μM A22 also had no apparent effect on cell morphology (Fig. S7A). Both A22 and MP265 were active in BSK-II medium as 50 μM of either drug induced cell rounding in BSK-II-grown cells of the *E. coli* strain MC1000 (Fig. S7B). We also measured the effects of A22 and MP265 on *B. burgdorferi* growth. The occurrence of cell growth is an important consideration, as cell rounding or bulging phenotypes associated with MreB inactivation are growth-dependent (1). A22 did not affect the growth of *B. burgdorferi* when used at 50 or 500 μM, nor did MP265 when used at 50 μM (Fig. S7C). While 500 μM MP265 slightly reduced the growth of our strain (Fig. S7C), a significant amount of growth (about four generations) still occurred during the treatment. Thus, the lack of cell morphology defects in response of A22 or MP265 treatment cannot be attributed to a growth arrest. Our results therefore indicate that *B. burgdorferi* appears to be resistant to chemical inhibition of MreB by A22 or MP265.

### SUPPLEMENTAL TABLES

**Table S1.**
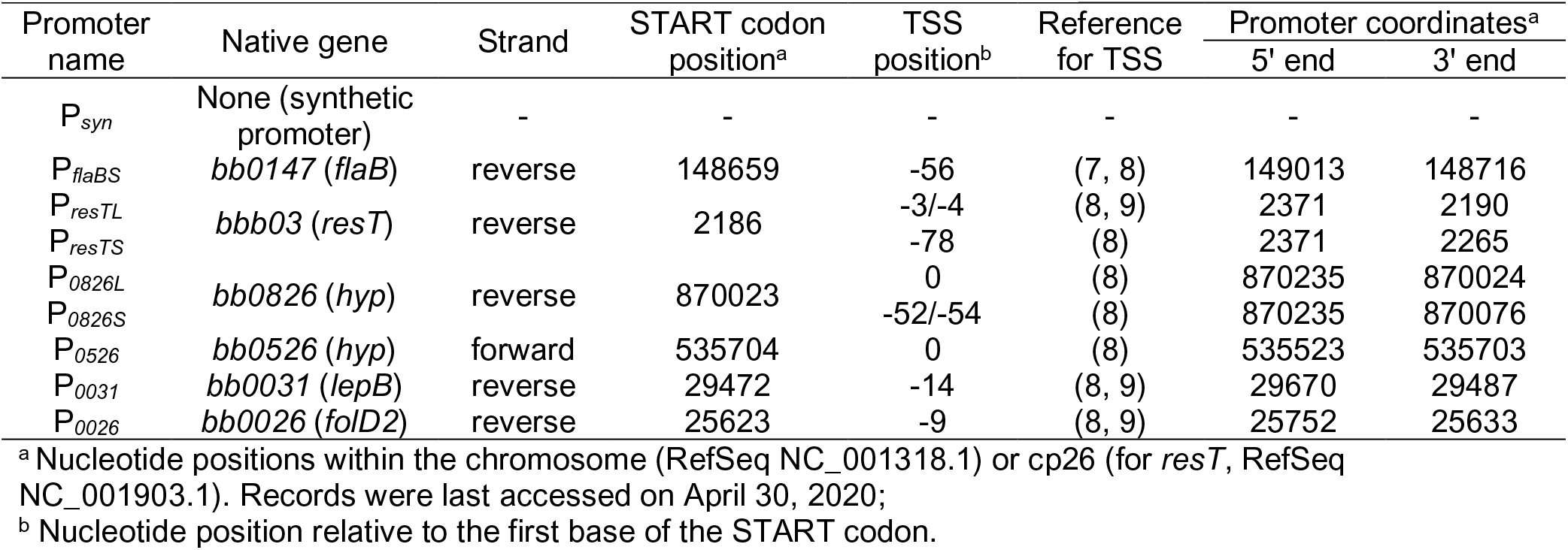
Promoters used to drive sgRNA expression in *B. burgdorferi*.

**Table S2.**
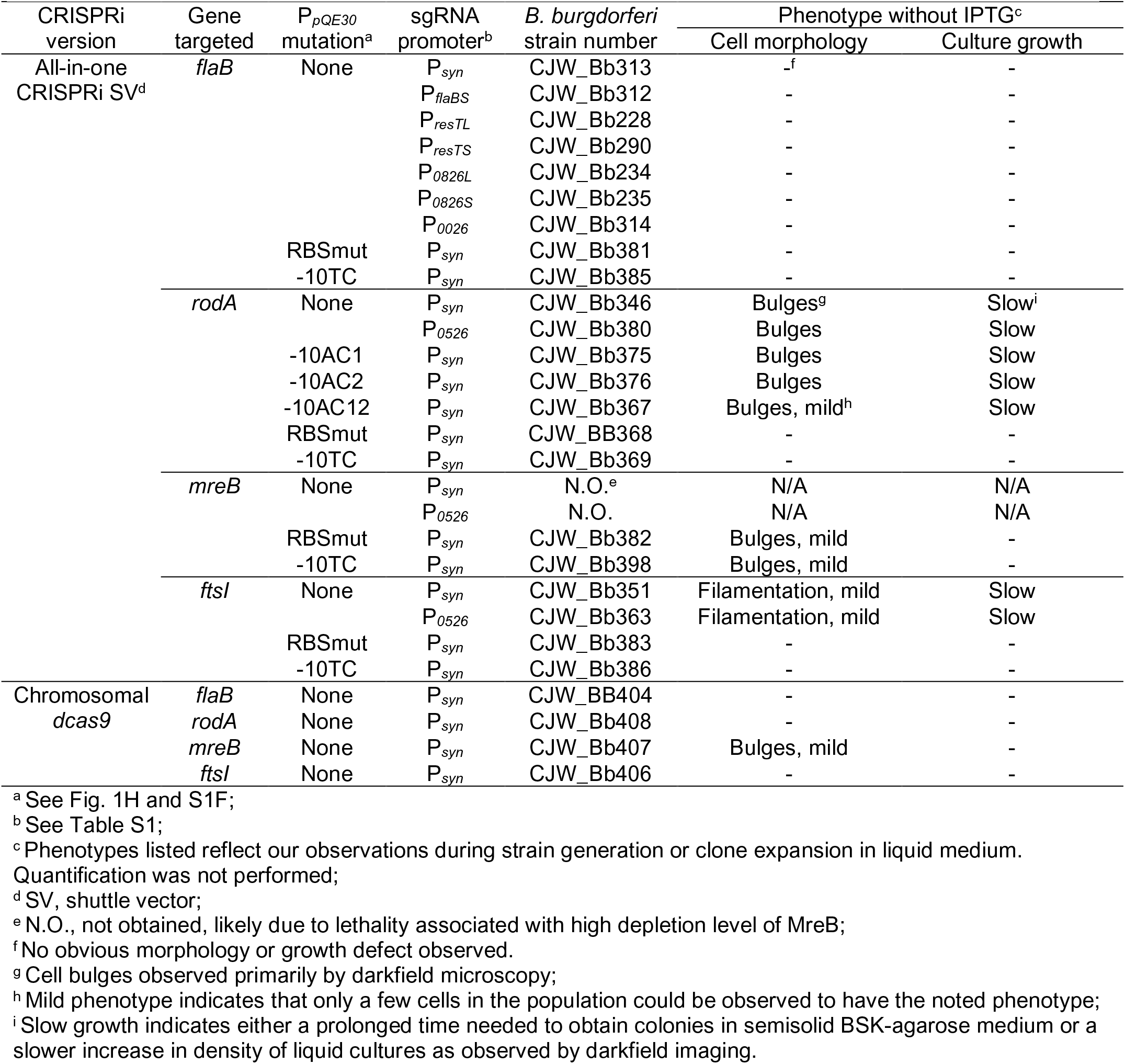
Morphology and growth phenotypes observed in the generated CRISPRi strains in the absence of IPTG induction of *dcas9* expression.

**Figure S1.**
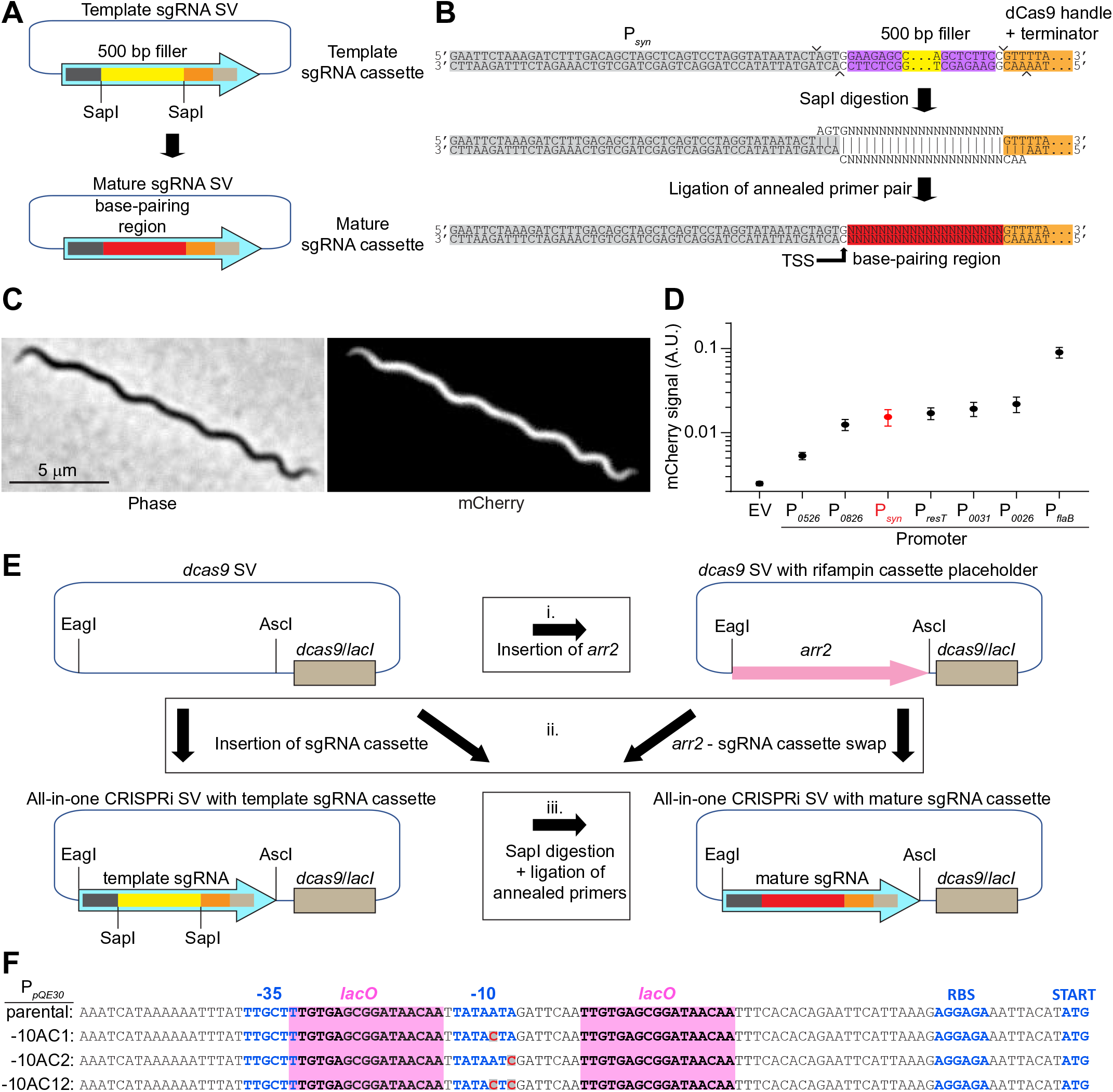
CRISPRi platform construction details. **A.** Schematic of conversion of a template sgRNA shuttle vector (SV, top) into a mature sgRNA cassette shuttle vector (bottom). **B.** Detail of the process outlined in A. The promoter shown is P*_syn_*. The 500 base-pair (bp) DNA filler is released by SapI digestion. Annealed primers are then ligated to the resulting backbone, generating the mature sgRNA’s base-pairing region placed immediately downstream of the promoter’s transcriptional start site (TSS). **C.** Phase-contrast and fluorescence images of a cell of strain CJW_Bb122 expressing mCherry from P*_syn_*. **D.** Quantification of the P*_syn_* strength through cellular mCherry fluorescence intensity measurements. The data for all other promoters was obtained and presented in reference (10). A.U., arbitrary units; EV, empty vector. **E.** Cloning avenues for generation of all-in-one CRISPRi shuttle vectors. Top left: *dcas9* shuttle vector. Insertion (cloning path i.) of a rifampin resistance cassette, *arr2*, between its AscI and EagI sites generates a *dcas9* shuttle vector with a rifampin cassette placeholder (top right). Insertion of a sgRNA cassette between the same sites of the *dcas9* shuttle vector (cloning path ii.) yields all-in-one shuttle vectors carrying either a template sgRNA cassette (bottom left) or a mature sgRNA cassette (bottom right). Using the *dcas9* shuttle vector with a rifampin cassette (top right) as a starting point in this cloning step offers the advantage of easy screening of clones by checking for loss or rifampin resistance. Conversion of the template sgRNA cassette of the all-in-one CRISPRi shuttle vector (bottom left) into the mature sgRNA cassette (bottom right) requires SapI digestion followed by ligation of annealed primers (cloning path iii.), as outlined in panel B. **F.** Mutations, other than those shown in Fig. 1H, introduced into the P*_pQE30_* sequence in an attempt to decrease basal expression of *dcas9*. *lacO*, LacI binding sites; RBS, ribosome-binding site.

**Figure S2.**
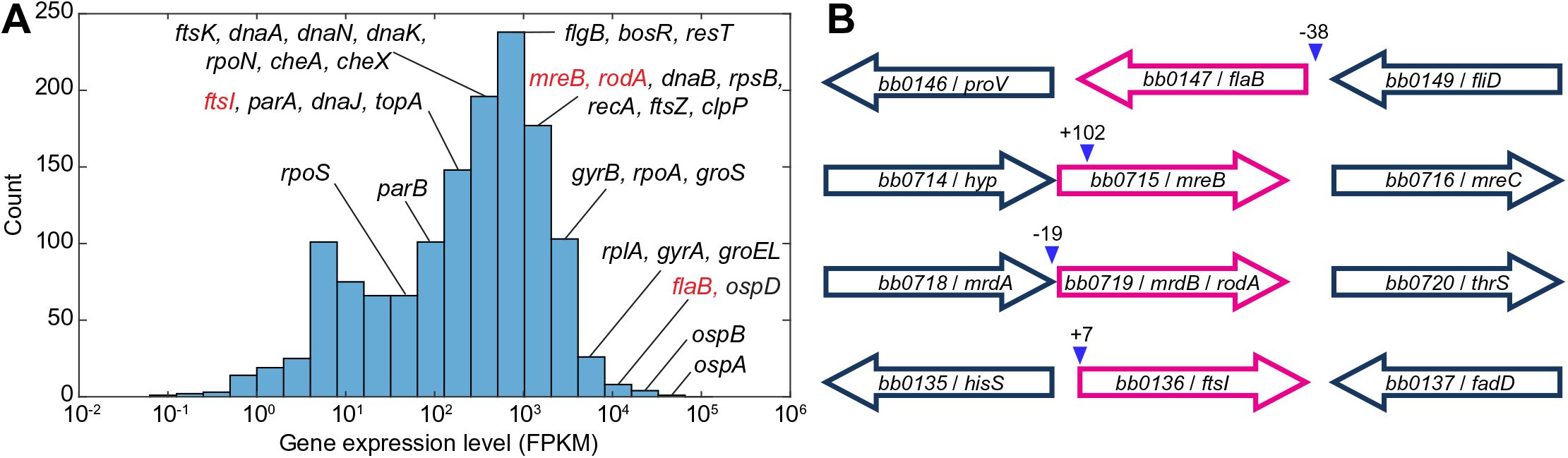
Genes targeted by CRISPRi in *B. burgdorferi*. **A.** Histogram showing the distribution of *B. burgdorferi* gene expression levels. The RNAseq data was obtained and reported by Arnold el al. (9) using strain B31-A3 (11) grown in BSK-II liquid culture. The data shown is from the early exponential condition as described in the original publication. Each bin represents a 2-fold range of expression levels. Indicated are expression bins where many physiologically important genes are located. Highlighted in red are the genes targeted by CRISPRi in our study. FPKM, fragments per kilobase per million reads. **B.** Genomic context of genes targeted by CRISPRi and location of CRISPR target sites. Coding regions of targeted genes are shown in pink, while upstream and downstream genes are in dark blue. When the distance between two adjacent genes is so short as to suggest that the genes form an operon, the genes are drawn close to each other. However, previous studies (7–9) have identified transcriptional start sites immediately upstream of the START codons of each of these genes. Labels contain both the gene number and its abbreviation. Blue arrowheads show the location of the CRISPR target sites, always within the 5′ UTR or the coding sequence of the gene. Values indicate the nucleotide position relative to the translational start site where the 5′ foremost complementary base of the sgRNA binds. Arrows are not drawn to scale.

**Figure S3.**
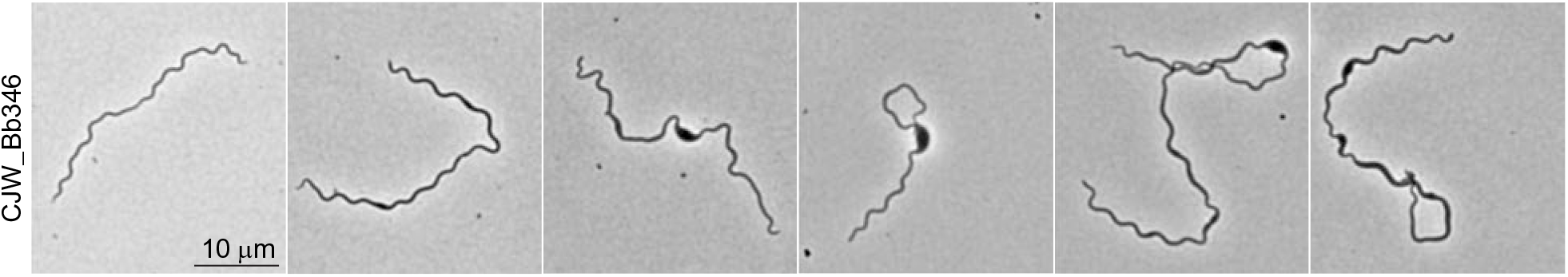
RodA depletion phenotype in the absence of IPTG induction. Phase contrast images of cells of strain CJW_Bb346 grown without IPTG. A cell with a normal width is at the left, while the other panels show various degrees of cell bulging.

**Figure S4.**
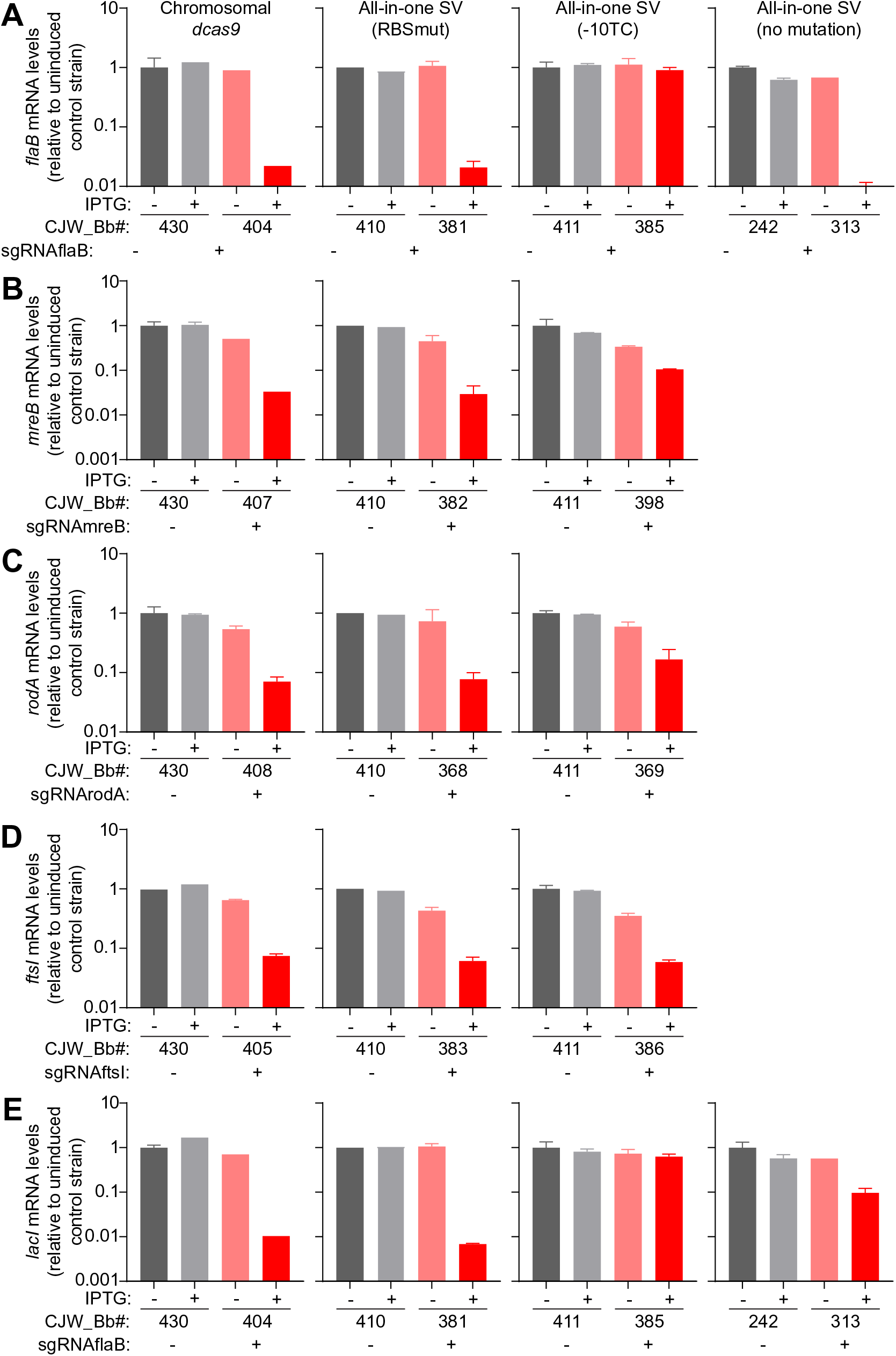
Effect of CRISPRi on targeted gene mRNA levels. **A.** *flaB*, **B.** *mreB*, **C.** *rodA*, **D.** *ftsI*, and **E.** *lacI* mRNA levels measured in the indicated control strains (gray) and CRISPRi depletion strains (pink and red) after two days of growth with or without IPTG. Shown are the means ± standard deviations measured from two cultures, or the values of single measurements (when no error bar is present). The version of the CRISPRi platform carried by each set of strains is indicated above the corresponding column of graphs. SV, shuttle vector.

**Figure S5.**
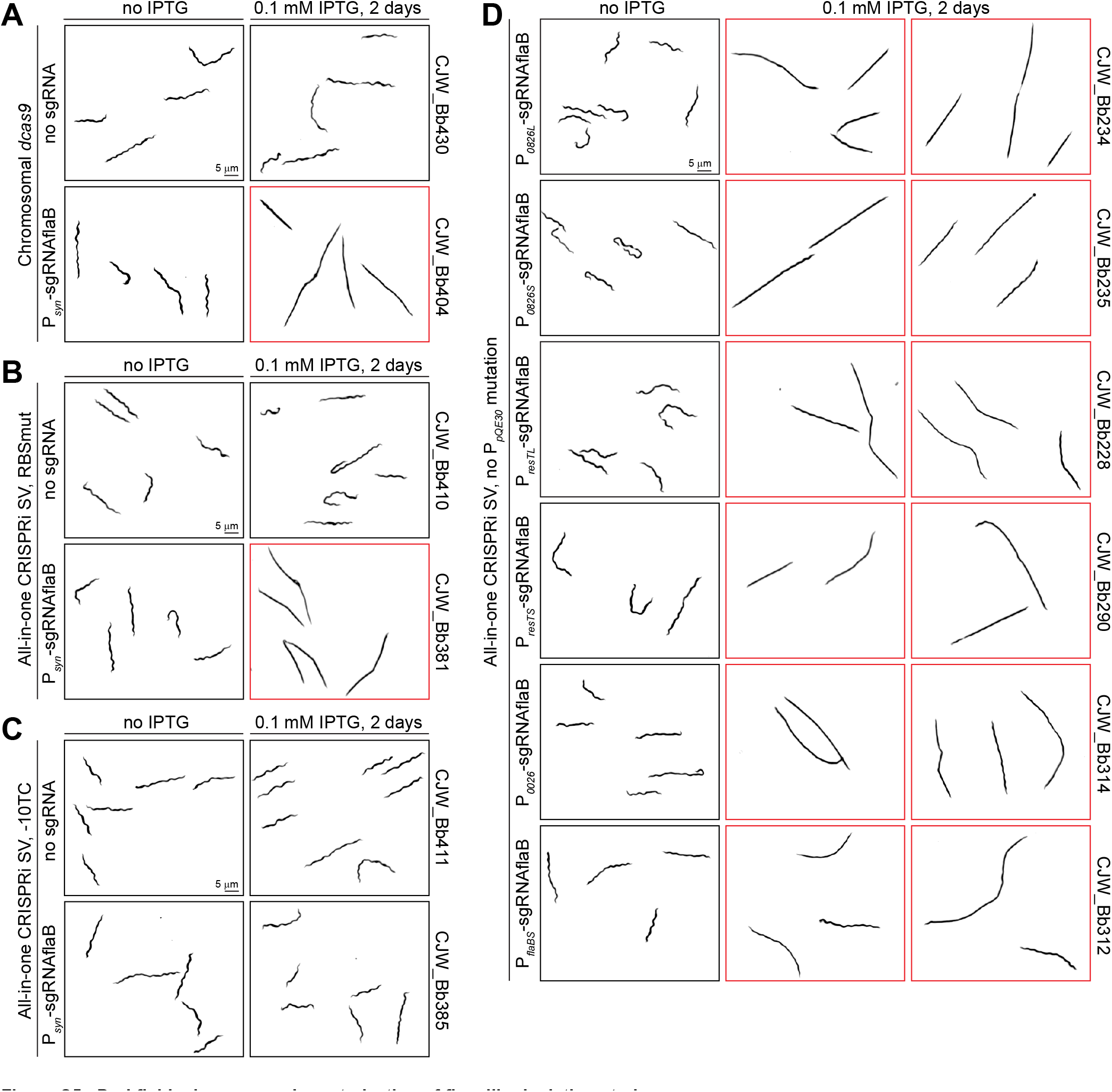
Darkfield microscopy characterization of flagellin depletion strains. **A-C.** Inverted darkfield images of strains expressing either no sgRNA or sgRNAflaB from versions of the CRISPRi platform that reduce the basal expression of *dcas9*. **D.** Inverted darkfield images of strains carrying *flaB*-targeting all-in-one CRISPRi shuttle vectors. The strains differ only in the promoter used to drive sgRNAflaB expression, as noted. **A-D.** Images showing a flagellin depletion phenotype are outlined in red. SV, shuttle vector.

**Figure S6.**
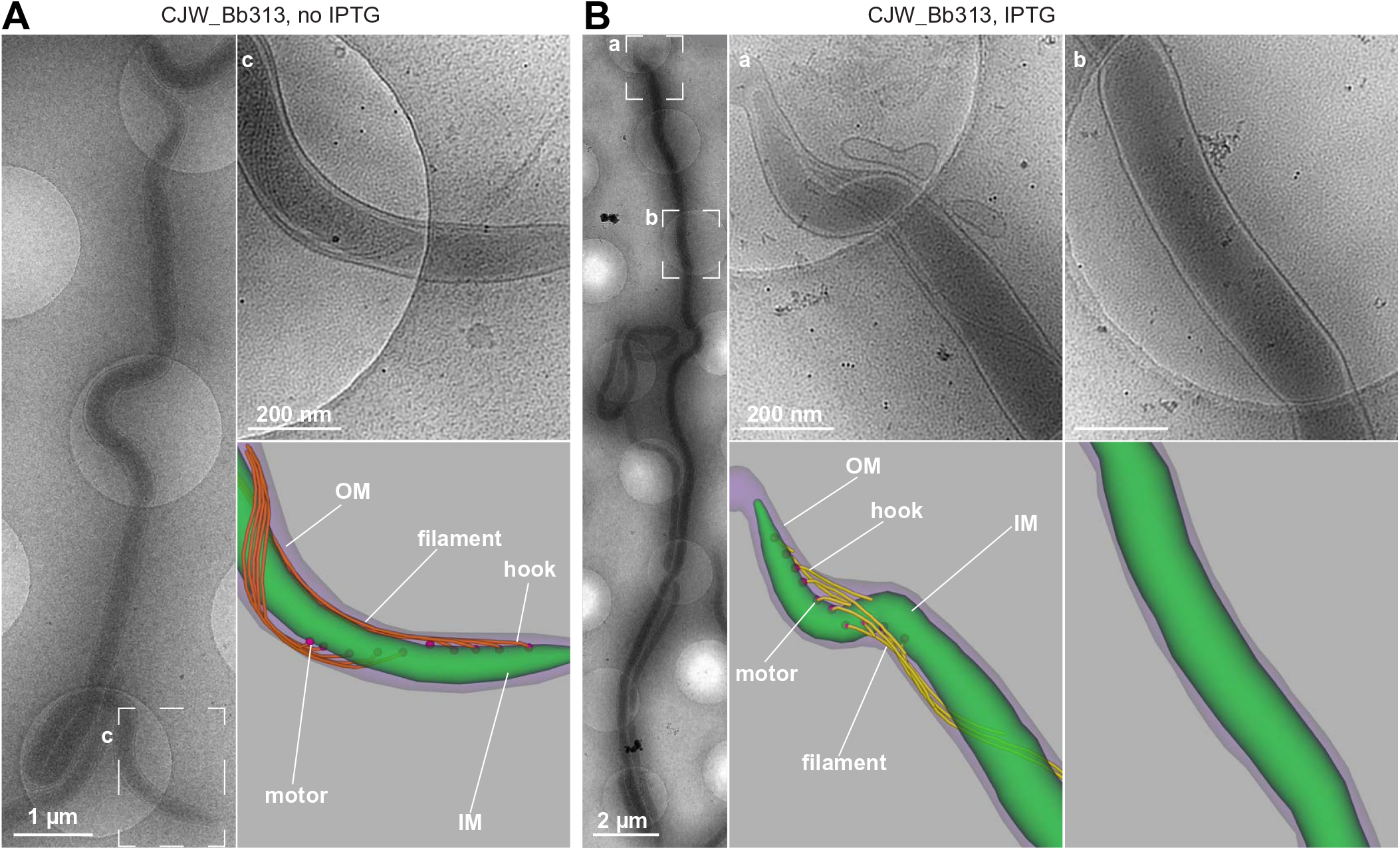
Cryo-ET characterization of flagellin depletion. **A.** Cryo-ET-based detection of periplasmic flagella in a cell of strain CJW_Bb313 grown in the absence of IPTG. Shown at the left is a low magnification view of the same cell as in Fig. 4C. Top right: high magnification view of the bottom end (c) of the cell. Bottom right: three-dimensional segmentation of the bottom end region of the cell. **B.** Flagellin depletion assessed by cryo-ET in a cell of strain CJW_Bb313 after two days of IPTG exposure. Left: low magnification view of the entire cell. Top center and right: high magnification views of the end (a) and center (b) of the cell, respectively. Bottom center and right: three-dimensional segmentation of the end and center of the cell, respectively.

**Figure S7.**
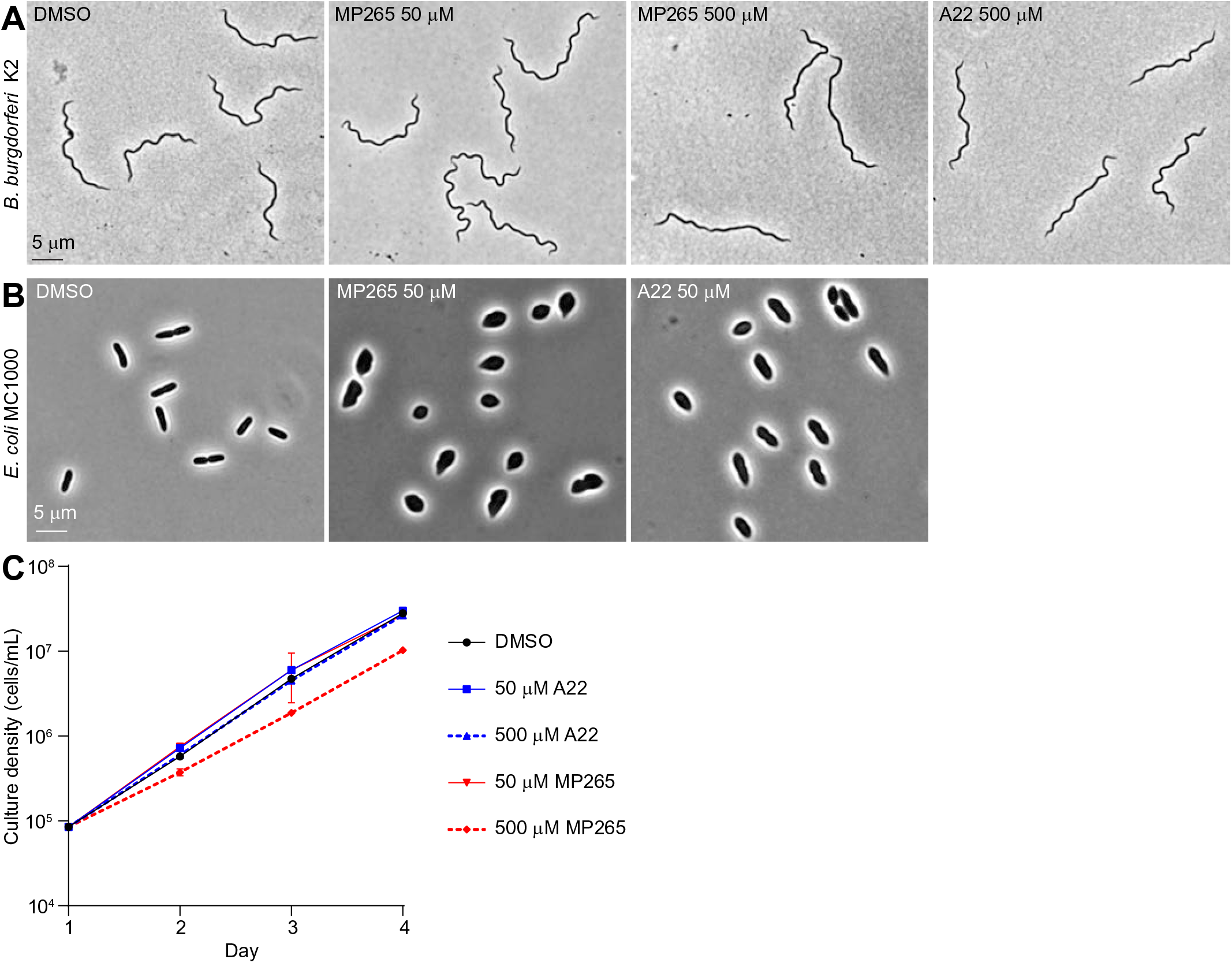
Phenotypic characterization of *B. burgdorferi* following exposure to MreB inhibitors. **A.** Phase contrast images of cells of *B. burgdorferi* strain K2 grown for two days in the presence of 0.1% DMSO or the indicated concentrations of MP265 or A22. **B.** Phase contrast images of *E. coli* strain MC1000 grown for one hour in BSK-II medium supplemented with DMSO, MP265, or A22. **C.** Growth curve of strain K2 treated with DMSO, MP265, or A22. Two replicate cultures were counted daily for each condition. Shown are means ± standard deviations.

### SUPPLEMENTAL MOVIE LEGENDS

All movies were recorded by imaging live *B. burgdorferi* cells in BSK-H medium using darkfield microscopy. Stream acquisition with an exposure time of 200 ms was used. Shown are inverted images. Elapsed time (in seconds) is displayed in the top right corner. A 10-μm scale bar is in the bottom right corner.

**Movie_S1.** A cell of strain CJW_Bb242 (control strain that carries a *dcas9* shuttle vector but does not express a sgRNA) grown in the absence of IPTG.

**Movie_S2.** Cells of strain CJW_Bb242 grown in the presence of 0.1 mM IPTG for two days.

**Movie_S3.** Cells of strain CJW_Bb313 (FlaB depletion strain carrying an all-in-one CRISPRi shuttle vector targeting *flaB* and the unmutated P*_pQE30_* promoter controlling *dcas9* expression) grown in the absence of IPTG.

**Movie_S4.** A cell of strain CJW_Bb313 grown with 0.1 mM IPTG for two days.

**Movie_S5.** Cell of strain CJW_Bb313 grown with 0.1 mM IPTG for two days.

**Movie_S6.** Cell of strain CJW_Bb313 grown with 0.1 mM IPTG for two days.

**Movie_S7.** Cells of strain CJW_Bb313 grown with 0.1 mM IPTG for two days.

**Movie_S8.** A cell of strain CJW_Bb313 grown with 0.1 mM IPTG for two days. The cell appears almost perfectly straight, like cells of a Δ*flaB* strain (12). The kink at the middle represents the division site.

